# Backpropagation-Based Recollection of Memories: Biological Plausibility and Computational Efficiency

**DOI:** 10.1101/2024.02.05.578854

**Authors:** Zied Ben Houidi

## Abstract

Since the advent of the neuron doctrine more than a century ago, information processing in the brain is widely believed to follow the forward pre to post-synaptic neurons direction. Challenging this view, we introduce the *backpropagation-based recollection* hypothesis as follows: *Cue-based memory recollection occurs when backpropagated Action Potentials (APs), originating in sparse neurons that uniquely activate in response to a specific trace being recalled (e.g. image of a cat), travel backwards. The resulting transient backpropagating currents follow the available open backward and lateral pathways, guided by synaptic weights or couplings. In doing so, they stimulate the same neurons that fired during the very first perception and subsequent encoding, effectively allowing a “replay” of the experience (e.g., recalling the image of the cat).* This process is pervasive, seen in tasks like cue-based attention, imagination, future episodic thinking, modality-specific language understanding, and naming.

After detailing our hypothesis, we challenge it against a thorough literature review, finding compelling evidence supporting our claims. We further found that gap junctions could be a plausible medium for such currents, and that cholinergic modulation, which is known to favour backpropagated APs and is crucial for memory, is a reasonable candidate trigger for the entire process. We then leverage computer simulations to demonstrate the computational efficiency of the backpropagation-based recollection principle in (i) reconstructing an image, backwards, starting from its forward-pass sparse activations and (ii) successfully naming an object with a comparable high accuracy as a state of the art machine learning classifier. Given the converging evidence and the hypothesis’s critical role in cognition, this paradigm shift warrants broader attention: it opens the way, among others, to novel interpretations of language acquisition and understanding, the interplay between memory encoding and retrieval, as well as reconciling the apparently opposed views between sparse coding and distributed representations, crucial for developing a theory of consciousness and the mind.

**Significance Statement:** Try to mentally picture the image of a cat. In this process, the word “cat” acted as a cue, and the fragile and non-persistent retrieved mental image is a recollected memory. Similar cue-based generative activities are ubiquitous in our lives, yet the underlying neural mechanisms are still a mystery. Neuroimaging and optogenetic-based studies suggest that cue-based recollection of memories involve the reactivation of the same neural ensembles which were active during perception (encoding). However, the exact neural mechanisms that mediate such reactivation remain unknown. We elaborate a novel hypothesis explaining how this can be implemented at single neurons: we hypothesize that the very same neural pathways used for perception are used backwards for recall, thus creating similar impressions during retrieval.

## 1 Introduction

Human brains learn to extract invariant representations from received sensory input in an unsupervised manner. However, mapping *signifiers* [1], i.e. mental representations of the image-sounds or “words”, to the *signified* mental representations that such words refer to, needs an interaction with an external agent that “supervises” the learning. Such “teacher” simultaneously generates the sound-image related to the signifier in the presence of the actual stimuli that relates to the signified: this can happen, as exemplified in Fig. 2, by generating the sound “cat” or the writing of the word “cat” in the presence of an actual image of a cat. When both are associated, we say that the agent has learned to *map the signifier to the signified*.

This familiar illustration is reminiscent of the general case of *cue-based recollection* of previously encountered stimuli. There, during *encoding*, the brain is exposed to the simultaneous *co-occurrence* of multiple stimuli. Later, during *recall*, one of these sensory stimuli, such as a visual scene or a smell, acts as a *cue* to trigger the retrieval of a related *memory trace* which co-occurred with the cue during previous encodings.

In this context, it is tempting to think at first that the repeated co-occurrence of both stimuli reinforces their connection, “in a hebbian manner”, thus allowing the mapping of cues to their related traces (and signifiers to their signified representations). However, it is commonly accepted that neurons process sensory information mainly in a forward manner, i.e. from pre-synaptic to post-synaptic neurons. Yet, in our problem, the two co-occurring signals (cue and trace or signifier and signified) would eventually reach a connection point after having followed only forward paths. Now, assuming, as discovered thanks to neuroimaging and optogenetic-based studies [2], that memories are stored and encoded in the same areas that uniquely responded to the memorized stimuli during the first encounter, a challenging question arises: What neural mechanism allows to associate one to the other, such that the activation of the signifier or cue can trigger *back* the activation of the signified or trace and vice versa.

In this paper, we hypothesize that weak and fast-fading backpropagating action potentials (APs) (from post to pre-synaptic neurons) whose strength is proportional to pre-synaptic “weights” are the medium by which previously encoded information is recollected. Such signals start in what we call *source pointer neurons* and travel all the way back, selectively reactivating on their path the neurons which uniquely responded to the stimuli during the first encounter, thus creating a similar experience to the first time. A pointer neuron, as we develop later is a neuron that specializes, thanks to past memorized co-occurrences, in selectively and invariantly ^1^ responding only to the retrieval cue (e.g. signifier) or its associated memories to be retrieved (e.g. signified). We refer to this as *backpropagation-based recollection* throughout the paper.

After stating our hypothesis in the general case (Sec. 2.1) and illustrating it through the toy example of (modality-specific) understanding and naming (Sec. 2.2.2), we discuss its biological plausibility by running it against a thorough review of literature (Sec. 3). We found a significant body of experimental results corroborating the plausibility of most of its assumptions (See anticipated summary in Table. 1). We find, for example, that fading-away APs backpropagating to the dendrites have been widely measured, shown to exhibit interesting properties: they are stronger when the post-synaptic neuron is firing and most of all, they can be controlled by (e.g. cholinergic) modulation so as to increase their strength or disinhibit them (Sec. 3.1). Their role, however, was hypothesized to be mainly regulatory. We further found evidence for reverse top-down reactivation patterns inline with our hypothesis (Sec. 3.2.2). However, following the traditional conception of forward propagation, they were attributed (wrongly, we argue) to backward recurrent feedback connections. Crucially, we found that gap junctions, which are unlike prior belief, widely present in the cortex could be the channels through which backpropagated currents flow.

**Table 1.**
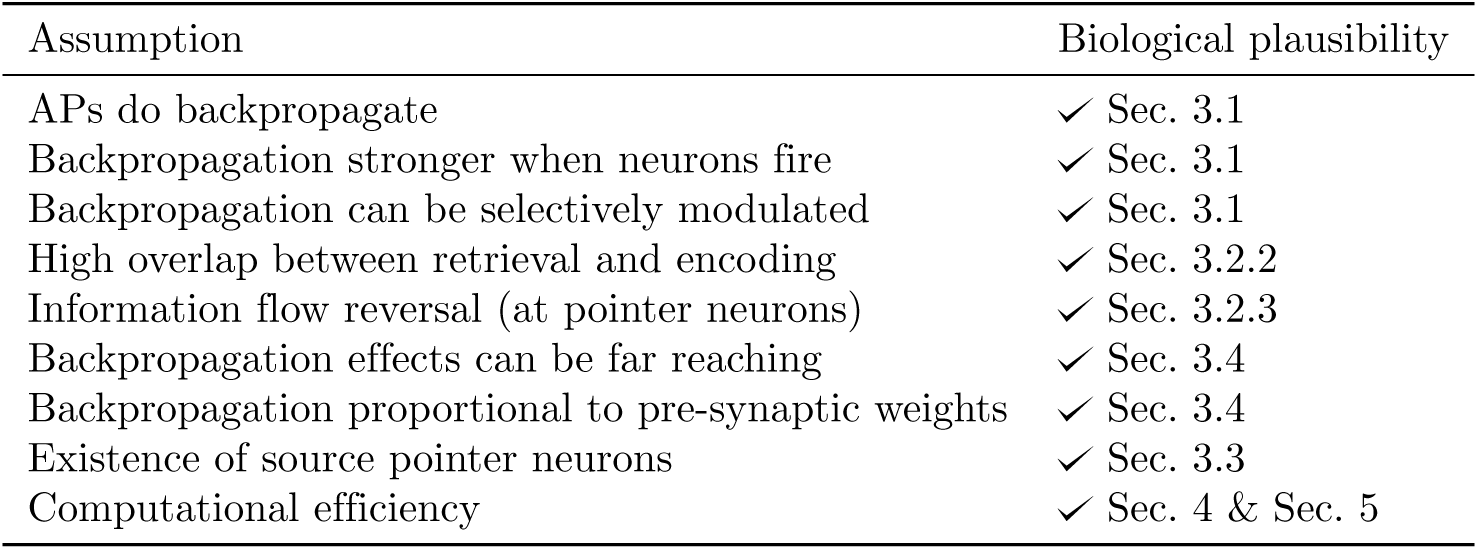
Summary of assumptions and supporting evidence (see Sec. 3.6)

| Assumption | Biological plausibility |
| --- | --- |
| APs do backpropagate | ✓ Sec. 3.1 |
| Backpropagation stronger when neurons fire | ✓ Sec. 3.1 |
| Backpropagation can be selectively modulated | ✓ Sec. 3.1 |
| High overlap between retrieval and encoding | ✓ Sec. 3.2.2 |
| Information flow reversal (at pointer neurons) | ✓ Sec. 3.2.3 |
| Backpropagation effects can be far reaching | ✓ Sec. 3.4 |
| Backpropagation proportional to pre-synaptic weights | ✓ Sec. 3.4 |
| Existence of source pointer neurons | ✓ Sec. 3.3 |
| Computational efficiency | ✓ Sec. 4 & Sec. 5 |

Finally, we demonstrate the computational efficiency of the backpropagation-based recollection principle leveraging (i) existing Spiking Neural Networks with *Spike Timing Dependent Plasticity* (STDP) learning [3, 4] as well as (ii) pre-trained neural networks from computer vision [5]. Through the first, we show how such mechanism can name an object as accurately as a machine learning classifier. On the second, we show how it is able to remarkably reconstruct an image starting from as few as a single sparse neuron, as anticipated in Fig 1.

**Fig 1.**
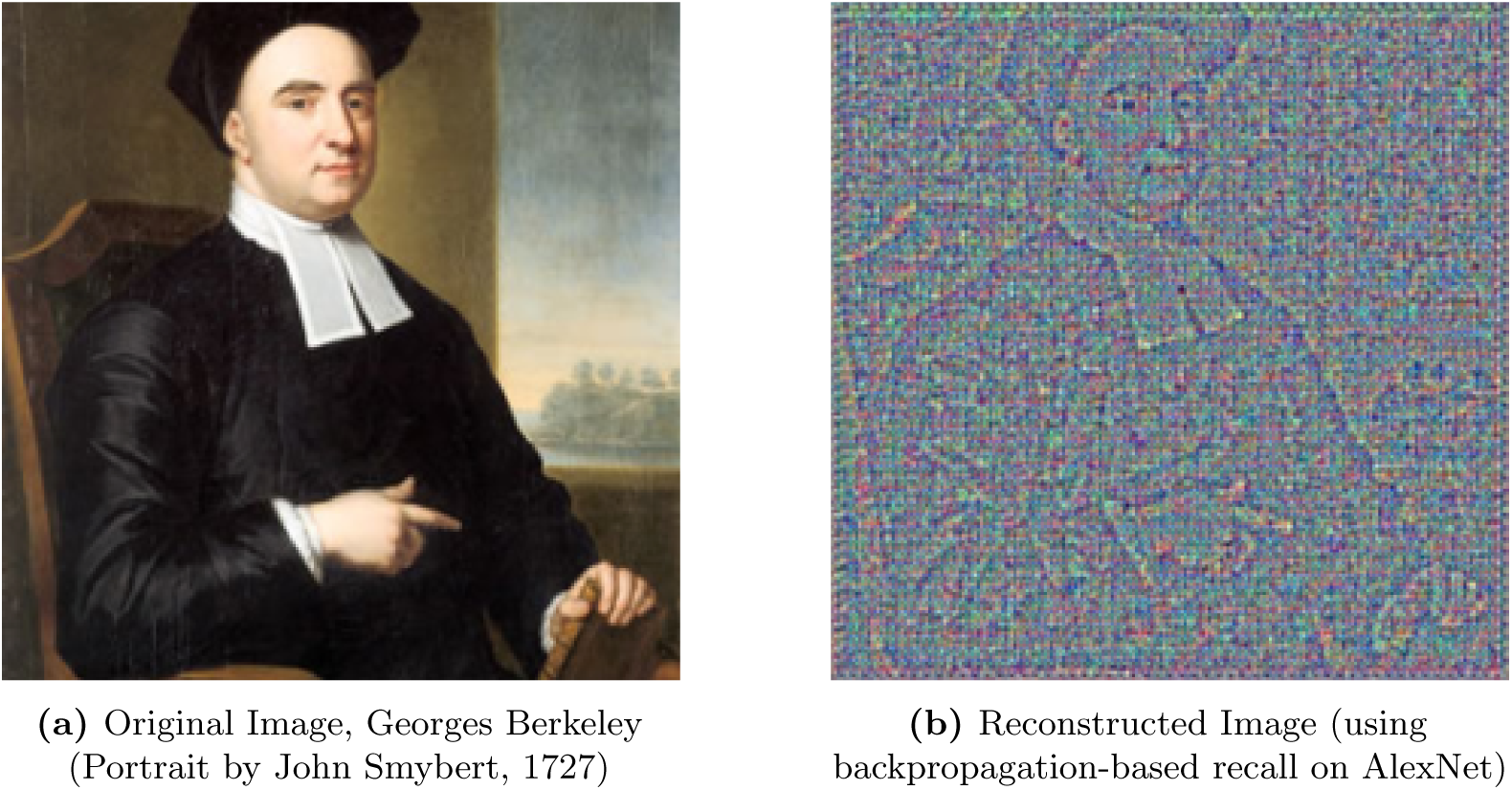
Comparison of original and reconstructed images.

**Fig 2.**
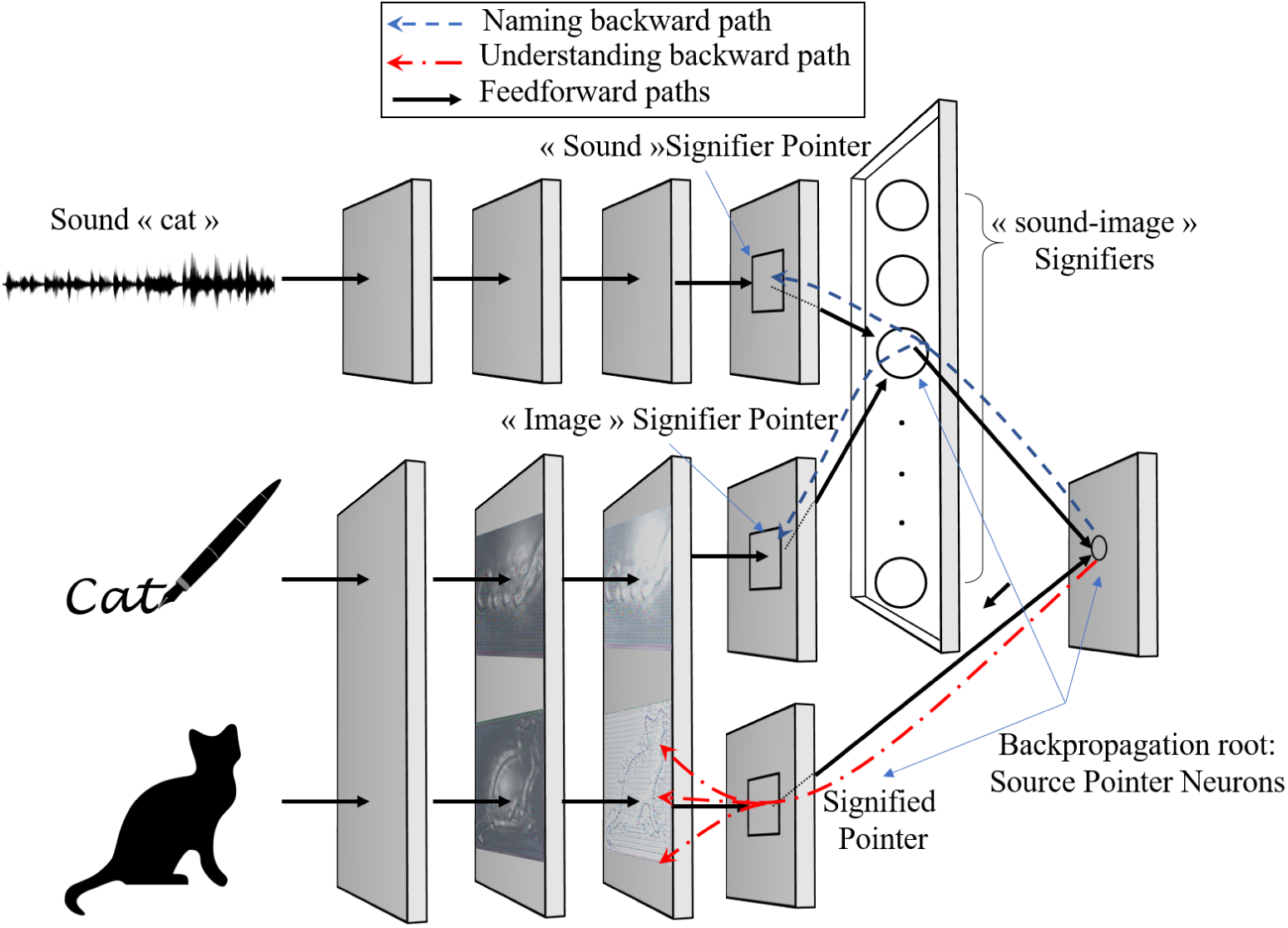
Schematized illustration in the case of naming and (modality-specific) understanding. Toy case where Boxes schematize consecutive processing stages from sensory input on.

This work is a considerable expansion of a preliminary version [6] of our research. The current paper adds more evidence of biological plausibility (e.g. role of ACh, gap junctions and mixed synapses), as well as the new study of image reconstruction capabilities.

## 2 The backpropagation-based recollection Hypothesis

### 2.1 General case

We posit our hypothesis and its assumptions as follows. *When a stimuli is presented to a (“trained”) brain, the latter responds by a unique activation pattern, resulting from the activity of an ensemble of neurons that signal the various salient features of the stimuli. Memories of past encountered stimuli are stored in a distributed fashion by the same neural ensembles that detected such stimuli when encountered and encoded for the first time. Recollection of memories is therefore a process by which the appropriate population of neurons is re-activated again so as to “re-live”, offline, an experience that is similar to the first encounter. We hypothesize that weak and fading-away backpropagating signals from post-synaptic to pre-synaptic neurons, whose strength is proportional to the post-synaptic neuron’s firing rate and pre-synaptic weights, is the mechanism by which the brain performs generative tasks. By generative tasks, we mean the regeneration of a previously lived and memorized stimuli (e.g. recollection in explicit declarative memory), the generation of a plausible future stimuli (e.g. future episodic memory) or the regeneration and combination of previous separately-lived and memorized stimuli, a process we refer to as imagination (e.g. imagining a cat that laughs by combining a memorized mental image of a cat with that of the act of laughing)*.

*The recollection process starts by the presentation of a retrieval cue, which activates few sparse neurons that uniquely identify both the retrieval cue and the to-be retrieved memory. We assume that these neurons learned to respond only to the presence of either of both stimuli thanks to a low-level learning rule such as STDP* [7]*; due to the (repeated or modulated) co-occurrence of both stimuli in the past: the cue and the to-be retrieved. We further hypothesize that the retrograde signal is initiated in such source “pointer” neurons (e.g. “Jennifer Anniston” cells* [8] *if the goal is to recall prior stimuli related to, say, Jennifer Anniston). The signal then travels first backwards (and then if the propagation path allows, also laterally) following the paths with high weights between coupled neurons, activating on its way all the various neurons that compose the mental images to be recalled. We hypothesize finally that such backpropagation can be controlled remotely via neuromodulation so as to invoke it, increase its strength or to inhibit it. This neuromodulation acts thus as a “switch” to control whether to retrieve or not*.

### 2.2 Naming and (modality-specific) understanding

Easier to illustrate our problem, a particular case of the above-mentioned cue-based tasks (which we partially simulate in Sec 4) is a form of explicit semantic memory related to *naming* objects and *modality-specific understanding*.

#### 2.2.1 Terminology

We build on the conceptualization introduced by swiss linguist Ferdinand de Saussure [1] when distinguishing between the *signifier* and the *signified*. We refer to the signifier as the mental representation of the sound-image of the word (i.e. the phonetic sound resulting from pronouncing it, or the image of its letters). We refer to the signified as the mental representation of the actual object that the sound-image refers to. Both the signifier and the signified are concepts, one represents the word, the other represents the mental image(s) that this word often refers to. Finally, we refer to *modality-specific understanding* as the act of mapping the signifier to its signified. *Naming* is instead *retrieving* the signifier that corresponds to a given stored mental representation or a presented live stimuli.

#### 2.2.2 Illustration

Fig. 2 illustrates our modeling through the example of three concurrent sensory inputs that are presented to a learner and that need to be “permanently associated”. This exemplifies a “teacher” that shows the “learner” an image of a cat, simultaneously to how the word cat is written, as well as the sound of the word. This can be also simply seen as three concurrent cues We assume as illustrated that there exists an area where the visual and sound activation pathways intersect (here, at the last “backpropagation root” layer). Such intersection areas are the root of backpropagating APs. During the “understanding” task, the retrieval cue is a word-related stimuli (e.g. sound “cat” or image of the word) and the retrieved memory is the signified representation (illustrated by the understanding backward pathway). During naming, the retrieval cue is the signified object and the retrieved memory is the name or signifier of the object (illustrated by the Naming backward pathway).

In more details, sensory input of *a* sound “cat” as well as *an* image of the word are processed through consecutive feed-forward neural layers. Similarly to what happens in primates’ visual cortex [9], neurons in the earlier layers have learned (thanks to local rules such as STDP) to respond to simple features and the deeper we go, the more selective the neurons become and the more they respond to more complex features. Hence at a later processing stage, there exists fewer “sparse” neurons that selectively respond only to the presence of the sound “cat”, we refer to such neurons as the “sound signifier” pointer neurons of the word cat. Similarly, we assume that there exists neurons that selectively respond to the “image” of the word: we refer to such neurons as the “Image signifier” pointer neurons. Finally, neurons that selectively respond to both are called the “sound-image signifier” or simply signifier pointer neurons. Similarly, the visual image of the cat itself is processed through various neural stages until certain neurons (referred to as signified in the figure) respond selectively only to the presence of the image of a cat.

This is how, as illustrated in the figure, the above described visual and sound pathways reach a common connection point, in a subsequent feedforward neural layer. We hypothesize that the repeated, or the neuromodulated^2^, co-occurrence of signifier and signified stimuli (i.e. saying “cat” in the presence of an actual cat) reinforces their connection, at this junction point layer, in “a hebbian manner” thanks to a simple rule such as STDP learning. Next, when presented with either signifier or signified related stimuli, the *source pointer neuron(s)*^3^ *in the backpropagation root layer will fire, resulting in backpropagating APs that are proportional to pre-synaptic weights*. The latter will cause the backward activation of the appropriate neurons, thus recalling the signified if the presented stimuli relates to the signifier, and vice versa.

It goes thus without saying that we adhere to the view that there exists few neurons that respond selectively to complex stimuli such as (i) sound signifier stimuli, (ii) image signifier stimuli, (iii) signified or (iv) uniquely to any of the three previous ones. During backpropagation-based retrieval, these neurons act as *pointers* to selectively reactivate an appropriate population of pre-synaptic neurons that uniquely characterizes the memory trace to be retrieved; thus creating a similar experience to that of the first encounter(s)^4^ when the memory was encoded. For example, what is retrieved could be an experience of the sound of the word “cat” with the particular voice or conditions in which it was encoded (in line with what is called the encoding specificity principle [10]).

This is how, as we will discuss later, *our hypothesis reconciles (i) sparse/localist and (ii) distributed representation theories in the brain, promising to end a long debate between cognitive psychologists and neuroscientists* [11, 12]. In our framework, there is no need to chose between them as both are needed, but for different purposes: Sparse coding, illustrated here by the presence of highly selective neurons, is needed for backpropagation-based retrieval, while the encoding of the entire memory trace is still done via a distributed set of neurons. The latter can be selectively reactivated “on-demand”, from source pointer neurons backwards. It is thus the simultaneous activation of an entire population of neurons that forms the entire memory trace, and single highly selective neurons are only pointers, helpful for retrieval.

Finally, and anecdotally, *the fact that backpropagating signals are “fading-away” in nature could explain the ephemeral nature of the experience of recalled memories or visual mental imagery’s lack of vividness*: the latter are not as persistent as the experience of live sensory stimulation.

### 2.3 What backpropagation-based recollection is not about

We stress that our hypothesis covers *only the recollection processes*. This means, in the particular case of explicit recall, the *reactivation, as close as possible, of the same neurons* that were activated during previous encounters. As a consequence, in the case of language and naming, what is covered is how to actually recollect the name, *not* yet how to *produce* it (e.g. emitting the sound or writing the letters). Further investigations are needed to reassess such “production” tasks in light of our new hypothesis.

Next, when we talk about understanding, we stress that we mean what is called the *modality-specific* features of semantic memory [13]: i.e. recalling how a face or an emotion looks or feels like exactly, as opposed to other aspects of semantic memory like finding abstract relationships between words. Worth mentioning though, Patterson et al. [13] reviewed semantic knowledge organization in the human brain and reported that all theories agreed on the fact that modality-specific recall is implemented by a distributed network, which is coherent with our theory.

Next, we assume that there are centers that remotely actuate via neuromodulation whether or not to call the recollection (by simply either increasing the backpropagation or inhibiting it). Our hypothesis does not cover what mechanisms control these control centers, and under which conditions recollection is favoured or shut down.

Finally, our hypothesis, being only focused on recollection, from sparse pointer neurons backwards, does not directly cover the mechanisms involved in sparse neurons formation, novelty or familiarity detection, and the interplay between short (e.g. few days back) and long (e.g. few years) term memories. Nonetheless, it still offers a novel ground to reason about these issues. *For example, the fact that the same sparse neurons keep being used to signal the presence of the same familiar stimuli throughout the years (e.g. face of own child) could explain why humans are unable to remember much younger versions of these faces, i.e. memories are updated in situ*.

## 3 Review of evidence in the literature

In this section, we position our hypothesis in the literature showing evidence that backs it up.

### 3.1 Existence of backpropagating action potentials

Despite the forward-processing prevalent view, a plethora of studies have measured, *in vitro* and *in vivo* in anesthetized [14] and awake [15, 16] mammalians, action potentials (APs) that backpropagate to apical and distal dendrites, and this for various classes of neurons [17, 18, 19, 20, 21, 22, 23, 24], including in hippocampal pyramidal neurons [22, 23, 24], an area that we suspect to be a “backpropagation root layer” (see Sec.3.2.3 later). We cite in the following some of them. For a more complete list, the reader can refer to the reviews of Stuart et al. [17] or Waters et al. [19] which also hypothesized few roles of backpropagated APs.

Williams and Stuart [20], for example, performed simultaneous somatic and dendritic recordings from Thalamocortical (TC) neurons and measured that APs, both due to sensory information or cortical excitatory post-synaptic neurons, backpropagate into the dendrites. The same authors [21] further measured the same phenomenon in neocortical pyramidal neurons. Interestingly for our hypothesis, they found that APs due to physiological patterns of firing, backpropagate three to four times more effectively compared to APs pertaining to mean firing rates. This observation was confirmed by several other studies (reviewed by Waters et al. [19]) which found that backpropagation was modulated by synaptic input, including in hippocampal pyramidal neurons [22, 23, 24] of interest to our hypothesis. For example, properly timed excitatory input leads to the amplification of backpropagation, whereas inhibitory input might block it.

Next, more interestingly for our hypothesis given the role of the hippocampus in initiating retrieval, it was shown that neuromodulation therein can further influence backpropagation, for example selectively and considerably enhancing it. Indeed, in hippocampal pyramidal neurons, many studies have observed that cholinergic agonists –which mimic the effects of acetylcholine– enhance the backpropagation progressively, leading to stronger and stronger effect on subsequent APs [25, 26, 27, 28].

*Neuromodulation can thus act as a “switch” to enable or disable AP backpropagation, a feature that is necessary for our hypothesis:* as described in Sec. 2, not every presentation of a visual stimuli systematically leads to the activation of “naming”, and not every presentation of signifier stimuli leads to the evocation of its signified representation: the brain has mechanisms by which it controls when to recall and when not.

*Finally, and perhaps equally intriguing, the very same acetylcholine (ACh) is widely documented to trigger bursts of synchronized neural activity initiating in the hippocampus*[29, 30] *which as we will argue later, in Sec. 3.4, could be the consequence of backpropagating currents flowing through gap junctions, to cause memory trace reactivation. It is also accepted, as we will document later in Sec. 3.5, that such cholinergic modulation is crucial for the hippocampus-mediated declarative memory, as opposed, e.g. to procedural memory* [31]

*Nonetheless, to the best of our knowledge, the role of such activity-dependent backpropagation of APs, as reviewed in the literature* [19]*, has been so far hypothesized to be local, acting as a feedback loop from post-synaptic to pre-synaptic neurons, to regulate spiking activity or support synaptic plasticity. In this paper, we suggest instead that their effects further backpropagate, playing a direct role in cue-based recall.*We next oppose our hypothesis to the state of knowledge in experimental cognitive sciences about explicit memory and the neural correlates thereof.

### 3.2 Explicit memory: neural correlates

Our hypothesis applies to any task in which the reconstruction of previously encoded stimuli is explicitly needed: we talk about explicit or declarative memory. The literature on this topic found its origins in early experimental cognitive science research (e.g. Tulving and Pearlstone [32], Tulving et al. [33]) before recent advances, driven by neuroimaging and optogenetic stimulation, shed more and more light on some actual neural correlates [2].

#### 3.2.1 Role of cues: encoding specificity and ecphory

##### In experimental cognitive science

One crucial principle in explicit memory was formulated by Tulving and Thomson under the name of encoding specificity [10]. The principle stresses the role of retrieval *cues* and the entire *context* that is perceived during encoding: elements of the surrounding context that was present during the first perception and encoding moment can act as efficient retrieval cues in the future. This process by which an internal or external sensory cue activates the stored memory trace is known as ecphory [34, 35]. Such early work [32, 10] played a role in differentiating between *availability* of memories and their *accessibility*, thanks to cues, let them be internal or external stimuli-based: the inability to recall does not necessarily mean that the memory is not available but could be also due to the lack (inactivation) of appropriate cues. Our hypothesis offers a neurobiological ground to interpret and simulate the above principles: for us, any surrounding context during encoding can act as a retrieval cue as long as it activates the source pointer neurons that uniquely^5^ identify the retrieval cue and all the surrounding context to be retrieved. These source pointer neurons activate back the appropriate pre-synaptic populations, resulting in the recall of all related traces.

*Following our hypothesis, we see how two components may affect memory retrieval as predicted by Tulving: the lack of the appropriate cue (no cue, no backpropagation) or a decay of synaptic weights (backpropagation cannot reactivate the trace). Our hypothesis further predicts two others: (i) a decline in the intensity or extent of the backpropagation, e.g. an impairment in the neuromodulation that facilitates backpropagation can hamper retrieval. Conversely, (ii) an excess of such backpropagation leads to higher levels of “intrusive thoughts”*.

Today, encoding specificity has passed the test of time [2]. Its “opponents” [36, 37, 38] do not question the necessity of some degree of match between encoding and retrieval conditions, but rather emphasize additional factors that influence retrieval performance; the most important being the “discriminative” power of the retrieval cue or its distinctiveness: recall performance is not only impacted by the degree of match between encoding and retrieval conditions, as thought earlier, but also by cue overload, or to what extent the cue is “discriminative”.

*Our framework natively predicts this effect. With respect to Fig.2, if the image of a cat (cue) appears, during learning, simultaneously with all sorts of names, and not only “cat”, then the backpropagating APs would simultaneously activate many “Signifier” neurons (and not only that of the cat), making it hard to distinguish and hence name. This necessity of being discriminative can be seen later in our simulations (Sec. 5.2): after backpropagating APs to the “categories” layer, the “signifier neuron” that gets the highest “votes” is elected to signal the name of the object. If all neurons receive “equal votes” because of cue overload, it would be impossible to retrieve the right name*.

##### Neural correlates

We next further challenge our hypothesis by diving into the neurobiological foundations of encoding specificity and explicit-memory, which saw tremendous advances driven by both neuroimaging and optogenetic stimulation. In a recent thorough review [2], a large body of research strongly supported Semon’s and Tulving’s cognitive theories: namely, that (i) the success of accessibility depends on the interaction between cues and memory traces and that (ii) there are strong ties between encoding and retrieval, including at the level of activated neural ensembles. In particular, it was shown, using artificial optogenetic stimulation ^6^ that it is possible to mimic (or disrupt) ecphoric processes by activating (or inhibiting) the *same specific neural ensembles* that were active during encoding^7^. In an experimental study [39], blocking the neural ensembles that were used to recognize the cue during encoding, resulted in retrieval impairment: Mice were conditioned to produce a fear response whenever placed in a particular context (i.e. the cue). At the same time, CA1 neural ensembles that were particularly active during encoding are optogenetically tagged. Whenever placed in the same context again, mice expectedly freeze as a sign that they recognize the environment. However, placing them while inhibiting the previously tagged neural ensembles considerably reduces freezing levels. *This means that if the neural ensembles that recognize the cue do not activate, the memory is not retrieved.* Other studies (e.g. Liu et al. [40]) successfully demonstrated the complementary direction: artificially reactivating the neural ensembles that recognize the cue induces the retrieval of the memory trace (i.e. freezing), even outside the context in which the conditioning happened.

In the previous two families of experiments, (i) mapping cues and traces was learned naturally, (ii) neural ensembles were tagged depending on their activity during encoding, and (iii) artificial inhibition or excitation was used to disrupt or elicit retrieval. A recent family of experiments [41] showed that it is even possible to associate cues and traces artificially and to later elicit retrieval in natural conditions. In particular, they repeatedly used photo-stimulation to artificially activate a neural ensemble that usually recognizes a special smell, simultaneously to photo-stimulating another that elicits avoidance. After this co-occurrence-based conditioning happened, mice exposure to the real smell led to avoidance: mice “remembered” to avoid although they never experienced the smell in real conditions.

In optogenetic stimulation studies above, light selectively activates a set of neurons that were prepared in advance to become light sensitive. This allowed scientists to dissect the interactions between memory traces, retrieval cues and encoding. However, it is not clear today what mechanism in the brain can cause a similar selective reactivation, equivalent to optogenetic stimulation. *That’s exactly the purpose of our hypothesis: hence, our answer to the question “how does a neural ensemble activates in a selective way” lies in backpropagating APs following the paths with highest pre-synaptic weights. Our hypothesis also offers a framework to simulate (as we do later) and understand these issues at the level of single neurons*.

Summarizing, beyond confirming encoding specificity, the studies above [39, 40, 41] and many others in the thorough review of Frankland et al. [2] suggest that retrieval reactivates what was active during encoding, in a process often called *neural reinstatement*. *This principle is at the heart of our hypothesis as backpropagated APs should follow exactly the reverse path that uniquely led to the activation of source neurons during encoding. It turns out that many arguments support this reinstatement principle. We review them in what follows*.

#### 3.2.2 Retrieval as top-down re-activation

A significant body of research intentionally studied the overlap between encoding and retrieval and provided large evidence for top-down reinstatement using various ensemble tagging[40, 39, 42, 43, 44, 45, 46, 47, 48, 49], EEG[50] and neuroimaging[51, 52, 53, 54, 55, 56, 57, 58, 59, 60, 61] techniques. Furthermore, it was shown that the reactivation overlap between encoding and retrieval influences also the perceived retrieval quality [62, 63, 64, 58, 57]. For example, in the case of visual imagery (i.e. attempting to mentally visualize an image), it has been shown that activation overlap in visual cortex increased visual imagery vividness, or the subjective intensity of the remembered image[64, 58, 57]. We refer the reader to the references above and only discuss few examples of each technique.

In terms of neuroimaging, Dijkstra et al. [61] used Dynamical Causal Modeling (DCM) [65] to infer coupling between cortical regions involved in the tasks of visual perception as opposed to visual imagery. They measured that visual imagery vividness correlated more with top-down connectivity patterns (from high level cortical areas to lower level areas) as opposed to perception itself. Many other studies suggest such top down mechanism during visual imagery [66, 67, 68, 69, 70]. The reader can refer to Pearson’s recent review [70] of the cognitive neuroscience of visual mental imagery for more details about the top-down reverse hierarchy of information and the fact that the process seems to be a weak form of the bottom-up perception. *However, due to the widespread view that neural computation is mainly forward from pre-to-post-synaptic neurons, this top-down activation cascade has been often interpreted as the result of feedforward feedback connections from higher-level cortical layers to lower ones*.

Then, and perhaps more convincing than neuroimaging, neural ensemble tagging techniques also confirm the principle. In addition to work we described above [39, 40, 41], another recent example is Guskjolen et al. [49] who performed contextual fear conditioning experiments on young mice while tagging the neural ensembles which were active during encoding. As happens with infantile amnesia in humans, the infant mice later exhibited forgetfulness. However, photo stimulation of tagged neurons, only in the hippocampal formation (Dentate Gyrus in particular), induced memory recovery and reactivation of broader areas which were tagged during conditioning including hippocampal CA1 and C3, and cortical neurons. Beyond supporting neural reinstatement, this finding also supports the idea that traces are distributed in neural ensembles that span many cortical areas [71], each responsible of one aspect of information (sensory, motor, visual etc).

#### 3.2.3 Existence of “backpropagation root layers”

Guskjolen et al. [49] leads us to the last point we review in this section: the role of the Medial Temporal Lobe and its relationship to our *hypothesized backpropagation root layers, where our source pointer neurons would be located. If correct, this area should form the “glue” between cues and retrieved traces, and should observe a reversal of information flow. Interestingly, the Medial temporal lobe and the hippocampus in particular have been shown to (i) play this role and (ii) exhibit a similar reversal behavior*.

For (i), many theories [72, 73, 74, 75, 76] claim that the hippocampus reinstates activity patterns in the cortex, which were alive during encoding. Studies like Tanaka et al. [39] above contributed supporting experimental evidence. As a reminder, by permanently tagging neurons which were active during encoding (fear conditioning experiment where cue is the environment and trace is “fear”), they were able to later silence them with laser stimulation. Then, when selectively silencing only the tagged hippocampal CA1 cells (as opposed to the trace ensembles), memory retrieval was impaired; and the rest of the neural ensembles in the cortex and amygdala (which encoded the fear response and used to reactivate during retrieval) were not reactivated again. Many other experiments [77, 78, 79] further show that retrieval success depends on whether or not the hippocampus was concurrently solicited (during both encoding or retrieval). Horner et al. [78] showed further evidence that the hippocampus binds together all elements composing a trace that is stored in distributed regions in the cortex, playing as a hub to perform what is also called pattern completion task. Finally, interestingly for our hypothesis, Staresina et al. [79] observed a reversible signal flow from the cue region to the “to be recalled” target region, through the hippocampus.

*This puts the HC and MTL in a good position to be our root backpropagation layers; implementing the link between cues’ networks and traces’ networks and being the place where flow reversal happens*.

Finally, citing verbatim a review and perspective from Moscovitch [80]: “Retrieval occurs when an external or internally generated cue triggers the hippocampal index, which in turn activates the entire neocortical ensemble associated with it. In this way, we recover not only the content of an event but the consciousness that accompanied our experience of it”. Referring to the hippocampal memory indexing theory of Teyler and DiScenna [74], Teyler and Rudy [73], he adds: “Memory traces in the HC/MTL are encoded in sparse, distributed representations that act as an index or pointers to the neocortical ensembles that mediate the attended information”. *This claim, well-aligned with our source-pointer-neurons, leads to our next assumption which we further review*.

### 3.3 Existence of highly selective source Pointer Neurons

We assume that at a deeper level of processing, certain neurons become highly selective and invariant, serving as pointers to reconstruct the encoded stimuli they represent. For example, in the schematized case of language, some neurons would respond only to the signified or the signifier (one of the cues), and some would respond to both (i.e. what is in common between the cue and the “to be recalled”). Here, we scan the literature about concept representation, assessing our hypothesis’ plausibility. *It turns out that similar neurons have been documented and that their role is not yet well understood, given the still ongoing debate [12] between two opposed views: “distributed representations” [81, 82] and “ sparse coding” [83, 84]*.

Indeed, the distributed representations view defends that concepts are represented by the unique activation patterns of large populations of neurons. It is thus the pattern uniqueness across a large population that defines complex concepts, not particular single neurons. The sparse coding view defends instead that there exists few neurons that represent selectively particular items or concepts. The extreme version of sparseness would be that there is a unique cell that responds and represents each unique concept, what is pejoratively known as the grandmother-cell hypothesis [11, 12]. *Our framework reconciles both: information is stored in distributed networks, and sparse neurons, which also exist, play the role of hubs to connect such distributed networks, easing retrieval by being sources of backpropagated APs*.

In the literature, the first accounts of sparseness are old. Since the seminal work of Hubel and Wiesel [85], it became mainstream that neurons tend to respond to more and more complex features, the deeper we go in the processing layers of sensory input[9, 86]. Evidence suggests the existence of a hierarchy starting from the primary visual cortex V1, where basic features are encoded, until the inferior temporal (IT) cortex, where neurons selectively respond to complex shapes like hands and faces [87, 88, 89].

Other known examples of sparse coding, for spatial representation, are place and grid cells [90]. Place cells are neurons which signal specific places: as the individual navigates in its environment, only neurons that signal the current place field fire. Interestingly, such highly selective neurons were found within the hippocampal formation, our candidate backpropagation root layer as we discussed in Sec. 3.2.3.

Further, the literature is rife with studies that measured such selective neurons in the same MTL areas. For example, Fried et al. [91] have measured neurons that selectively discriminate humans (faces) from inanimate objects, interestingly during both encoding and retrieval. Others distinguished specific facial expressions. A little later, Kreiman et al. [92] have measured neurons that highly responded only to specific categories such as animals, houses and celebrities.

A continuous line of work [8, 93, 94, 95] has set to understand how the simple visual features are passed to upper layers of the hierarchy and how they are later used by higher cognitive processes: *a question to which our hypothesis is a candidate answer.* Quiroga et al. [8] reported the existence of highly selective neurons that responded to the presence of specific stimuli related to places or individuals such as Bill Clinton and Jennifer Anniston. One of the reported neurons even exhibited highly selective responses to any Halle Berry related stimuli, let it be the face or the written name. *The latter exhibits strikingly similar properties to our source pointer neurons as we schematized in language understanding and naming*.

Waydo et al. [94] later used a probabilistic approach to explore more rigorously original findings of Quiroga et al. [8]. Indeed, the latter obviously did not test all MTL neurons and all possible categories of objects. Hence, (i) a found selective neuron could respond to other untested categories and (ii) there might exist many neurons, and not only one as found by the authors, that would selectively respond to the same stimulus. Authors thus developed a probabilistic model to estimate the odds, confirming the sparseness hypothesis (yet, arguing also against single grandmother cells [83]). Nonetheless, the model leads to, as the authors conclude, only a bound on the true sparseness: the neural coding could be in reality even much sparser than they estimated.

Following-up, Quiroga et al. [93] further stressed that sparseness does not mean grandmother-cells, arguing against the unlikely possibility that only a unique neuron responds to each stimulus. *Here, although, for simplicity, we exemplify (Sec. 2.2) and simulate (Sec. 4) our hypothesis using single neurons, we do not assume their uniqueness. On the contrary, we believe multiple neurons with the same behavior are needed to guarantee resiliency in case of “failure”. Quiroga et al.* [93] *nonetheless conclude with a set of open questions, to which our hypothesis offers some candidate answers: “How are MTL cells involved in learning associations? How are MTL cells involved in free recall or the spontaneous emergence of recollection in the human mind?” We propose that backpropagated APs originating in these cells cause the recollection of related traces. Through neuromodulation, remote “control centers” act as a “switch” to facilitate recall or inhibit it. As we briefly discuss for mind wandering (Sec. 6.1), the same principle should apply to free recall*.

Finally, Quiroga et al. [95] have subsequently measured that MTL selective neurons reflect the subjects’ decisions about the stimuli rather than visual features themselves. To put this in evidence, they exposed subjects to confusing faces mixing two celebrities (e.g. presidents Bush and Clinton). As expected from previous studies, pre-exposing subjects to one celebrity, leads them to later “decide” that the morphed image pertain to the other celebrity. Later by recording Clinton’s and Bush’s neurons, they concluded that such MTL neurons fire in accordance to the subject’s decision. In an interesting follow-up comment, Reddy and Thorpe [96] remarked that damages to the MTL cause subjects to have memory impairments, yet keeping perfect perceptual awareness and consciousness. *This interestingly corroborates with our hypothesized role of source pointer neurons in triggering recall*.

*Summarizing, sparse neurons that respond selectively to complex concepts exist, in areas where we suspect them to do, with properties that are inline with our backpropagation-based recollection hypothesis*.

### 3.4 Gap junctions: a plausible conduit for Backpropagated APs

In our proposed model, identified cues will create a unique activation pattern, particularly triggering specific sparse neurons—likely within the MTL and the hippocampal formation—that are uniquely associated with those cues (as discussed in Sec. 3.2.3 above). The subsequent decision to evoke associated memories would be triggered by neuromodulators, like Acetylcholine which was shown to further promote backpropagating action potentials (see Sec. 3.5 below). These retrograde currents first return to the dendrites and then cascade backwards, and whenever possible laterally, aiming to access the broader neural network that encodes the relevant memory trace. To facilitate such a non-standard cross-cell flow of information, an efficient bidirectional communication system within the brain’s networks becomes paramount. Here, we delve into the potential of gap junctions in enabling such hypothesized retrograde currents. Traditionally recognized for their function in electrical synapses, gap junctions facilitate a rapid, bidirectional flow of ions and small molecules between adjacent cells. Beyond neurons, they are also present in glial cells, where they perform various functions that are currently being darstically re-evaluated [97, 98, 99].

For our hypothesis to stand, several criteria regarding gap junctions^8^ must be met: (i) Their presence should be significant, especially within the hippocampal formation and the neocortex; (ii) They should exhibit plasticity and be able to coexist harmoniously, sometimes simultaneously, with chemical synapses; (iii) The disruption or removal of these junctions should impede memory recall (iv) If Acetylcholine indeed instigates this recall process, its inhibition should also deter memory retrieval. Our literature survey has unveiled compelling evidence for all of the above, in addition to intriguingly elucidating the crucial role of both ACh, electrical synapses and gap junctions more generally, in triggering and driving synchronized neural activities across selected and distributed neural populations—vital for declarative memories.

In the remainder, while our primary focus (of illustrating biological plausibility of our hypothesis) will be on electrical synapses, we acknowledge that the other types of gap junctions may also play, perhaps a complementary, a part, as we will discuss below. We identify this as an intriguing avenue for future detailed investigation.

#### 3.4.1 Abundant prevalance against previous beliefs

Thinking about bi-directional inter-neuron communication, electrical synapses, once deemed peripheral in the central narrative of neuroscience, have been undergoing a renaissance the last 25 years. As defended by many recent reviews [100, 101, 102], despite revelations from the 1960s to the 1990s demonstrating their widespread presence across the adult mammalian CNS [103, 104, 105], electrical synapses were repeatedly unjustly attributed to limited regions, like the olfactory bulb. Historically, they were considered evolutionarily rudimentary—predominantly associated with invertebrates and ectothermic vertebrates—and were somewhat eclipsed by the more intricate chemical synapses [104]. Further, gap junctions, facilitating electrical synapses, were for a while thought to disappear with age and primarily function in developmental phases [106, 107].

However, as defended at length by Nagy et al. [100] and Pereda [101], their absence was only due to the difficultly of observing them, because of their tiny size and the technical requirements for proper tissue preparation and visualization. Further, since they were believed evolutionarily primitive and less complex, less efforts were spent to verify their presence and behavior in areas where they could perform complex computations as their more studied chemical counterparts.

In practice, the true potential of electrical synapses began to gain clarity with the discovery in 1998 of connexin36 (Cx36) [108], a new gap junction gene, that was the first connexin^9^ proven to be expressed in many neuron types in retina, brain and spinal cord [109, 110, 104, 102]. It should be noted that other types of gap junctions, mediated by other connexins, such as those found in glial cells and astrocytes, may also have untapped potential, although that is beyond the scope of our current investigation. Building on this discovery, antibodies tailored to Cx36 were quickly produced, providing a unique tool to study them. When utilized in immunofluorescence, these antibodies accurately pinpointed the location of Cx36 specifically within gap junctions [109, 111, 112], ruling out other potential intracellular sites [100]. This specificity, in addition to the analysis of structural properties, enabled precise discernment of Cx36-based neuronal gap junctions, finally proving their widespread presence across vast areas of the adult mammalian CNS [109, 111, 112, 113, 114, 115, 116].

In alignment with our hypothesis that memory recall can be initiated in the MTL/hippocampal formation^10^ but later retrieved from distributed cortical networks, electrical synapses have been widely found in the hippocampus [100, 117, 102, 118, 119, 113, 120, 121, 122] where their coupling mediates synchronous activity among inhibitory interneurons, and enables gamma oscillations in pyramidal cells. They were also found throughout the cerebral cortex[123, 124, 125, 126, 127] where they facilitate synchronous activities in both interneuronal networks and neocortical pyramidal cells [123, 124, 125, 126, 127]. As we will dive deeper in Sec. 3.4.3, such oscillations are interestingly disrupted in Cx36 null mice and when blocking gap junctions more generally [128, 129, 130, 131]. Moreover, these synapses are not limited to inhibitory interneurons but also exist among excitatory glutamatergic pyramidal cells of the hippocampus and the cortex [132, 133]. These projections are notably expansive, projecting into diverse regions of the brain, further providing a plausible ground for our hypothesis.

While their presence in other regions, such as the retina [134], olfactory bulb [112] among many others, may or may not directly relate our hypothesis, it nonetheless underlines their multi-faceted under-estimated roles that clearly go beyond the historically constrained narrative restricting them to developmental stages and specific neural circuits.

All in all, recent reviews argued that the relatively newly discovered ubiquity of gap junctions far exceeds our current understanding of their roles, suggesting there are significant undiscovered functions yet to be unveiled[100]. Their wide distribution —in both the hippocampus and other connected cortical areas—builds a foundational premise for our hypothesis. Now, given that backpropagated currents could theoretically flow between any pair of coupled cells—not just neurons—it raises the possibility that various other types of gap junctions might also be involved in this process. This idea merits deeper investigation in the future, especially in light of the evolving understanding and positive re-evaluation of the role of glia in computational and cognitive processes [97, 98, 135].

#### 3.4.2 Co-existence and “co-plasticity” with chemical synapses

To fully appreciate our hypothesized role of gap junctions in backpropagation-based recall, we must assess two pivotal points. First, electrical synapses (or gap junctions generally) and their chemical counterparts must co-exist, at least in source pointer neurons. Indeed, it is in such mixed-synaptic neurons that APs, ignited by forward-moving chemical synaptic activities, would reverse course via gap junctions during recall events. Second, and equally crucial, gap junctions must exhibit long term plasticity and this plasticity must align and adapt in tandem with the forward chemical pathway, so as to allow reverse tracing.

It turns out that since the discovery of their widespread presence, gap junctions have been consistently found to coexist with chemical synapses [113, 118, 114] and demonstrate intriguing plasticity behaviors[136]. Of high relevance to our hypothesis is the observation that both chemical activity and chemical neurotransmitters can modulate electrical coupling[137, 101]. This suggests that the learning observed in chemical synapses could be transferred to electrical synapses.

##### Existence of mixed synapses

Despite a history of under-reporting of the existence of gap junctions, more and more studies are reporting the co-existence of mixed chemical and electrical synapses in various areas of the brain, as reported by recent dedicated reviews [100, 138]. Among the ways to suspect or detect the existence of such mixed synapses, we find the joint immunofluorescence detection of connexin36 and vesicular glutamate transporter-1 (a marker for chemical synapses). In addition, mixed synapses can also be identified through advanced electron microscopy techniques [138]: Thin-section transmission electron microscopy (TS-TEM) reveals the structural details of gap junctions, while freeze-fracture transmission electron microscopy (FF-TEM) allows for the visualization of intra-membrane particles characteristic of these synapses. These methods, complemented by immunogold labeling, enable precise mapping and characterization of mixed synapses at the ultra-structural level.

Using such techniques, mixed synapses were reported in various areas in the mammalian brain [113, 114, 115, 116, 139, 120], and retina [109]. Closer to our theory, they were found in the hippocampus, as reported by many studies and reviews (e.g. [118, 119, 113, 120, 138])

As early as in 2000, Fukuda and Kosaka [118, 119] discovered that a subpopulation of hippocampal GABAergic neurons, containing a calcium-binding protein called parvalbumin, formed a network connected using simultaneously both types of synapses (Chemical contacts are located between axon terminals and somata, while gap junction ones are situated between the dendrites). Interestingly, the study revealed that this structure is not limited to the hippocampus but is also present in the neocortex, suggesting that it is fundamental structure of the cerebral cortex and concluding that this could be related to synchronized activities observed across various cortical areas, as would be suggested by our hypothesis.

Further evidence of existence in the hippocampus, beyond interneurons, was provided by Nagy [113] which found gap junctions at mossy fiber terminals (axons of granule cells that connect with CA3 pyramidal cells) and which have been known for having only chemical synapses. Beyond co-existence, electrophysiological studies have shown that mixed transmission actually co-existed between hippocampal mossy fibers and pyramidal cells[121]. Using thin-section electron microscopy, freeze-fracture replica immunogold labeling (FRIL), and dye-coupling methods, Hamzei-Sichani et al. [120] further confirmed the existence of mixed electrical-chemical synapses on both principal cells and interneurons in the adult rat hippocampus. The latter were found to be primarily glutamatergic. The synapses were found in several rat hippocampal regions, including, as in the previous work, CA3 pyramidal neurons and the dentate gyrus. Subsequent work on mixed neurotransmission in the mossy fibers of the hippocampus further argued that the phenomenon is likely prevalent in wide areas of the central nervous system (CNS), and that it is just a matter of time before this is properly documented [122].

More recently, noticing first that mixed synapses could lead to weird synaptic events (e.g. electrical currents preceding the chemical ones), Ixmatlahua et al. [140] interestingly revealed the unprecedented existence of gap junctions co-located with glutamatergic chemical synapses in axo-dendritic connections, and exhibiting a curious behavior where they are normally silent but could be activatable under certain conditions. This opens the way to more sophisticated mechanisms by which gap junctions are selectively used.

Finally, a more recent review[138] has provided an in-depth evaluation of the literature supporting the co-existence of mixed synapses, including growing, but needing further validation, evidence of mixed transmissions (beyond mere co-existence), leading the authors to wonder about the “enigmatic functions” of such mixed connections. Given the breadth of its cognitive functions, our theory is clearly an interesting candidate.

##### Plasticity and modulation by chemical synapses

Next, Despite a popular belief that electrical transmission is static and simple, more and more evidence reveals that it is much more dynamic, adaptive and complex than previously thought (see e.g. [141, 142] for an overview of such complexity). Over the years, electrical synapses have been reported to be highly plastic and capable of modifying their coupling strength under various physiological conditions [143, 144, 136, 145, 146].

For example, Haas et al. [144] observed an eLTP attributed to a coordinated burst firing between pairs of coupled Thalamic Reticular Nucleus (TRN) neurons. Wang et al. [146] later found that gap junctions in the same area undergo both long-term depression and long-term potentiation, which they termed eLTD and eLTP, analogously to plastic changes observed in chemical synapses. In particular, group I metabotropic glutamate receptors (mGluRs) induces LTD, while the activation of Group II mGluR, specifically mGluR3, leads to LTP. This indicates that different types of mGluRs can differentially modulate electrical synapses, hinting as well to a potential modulation of electrical synapses by chemical activity, which we’ll see in the next paragraph. Indeed, through the release of neurotransmitters, chemical synapses can activate various receptors (like mGluRs in this study) which can lead to intra-neuron signaling cascades that modify the strength of electrical synapses. Besides, APs generated at chemical synapses can propagate to regions of the neuron where gap junctions are located. Depending on the state of the electrical synapses (strengthened or weakened), this can influence the synchronization of activity between neurons. For example, if electrical synapses are potentiated, action potentials might more effectively spread to adjacent neurons, enhancing synchronous activity.

But beyond plasticity, what is interesting for our theory is the interplay between the chemical activity and the electrical one, and *in particular the ability of chemical coupling to influence also electrical coupling*, as this is needed for our theory (*so that the backpropagated currents, mediated by gap junctions, follow the same paths followed by chemically mediated forward currents*).

In this context, more and more evidence supports the notion that chemical coupling can influence and potentially enhance electrical coupling, a probably necessary feature for our theory. This is grounded in the observed modulation of gap junction proteins by (i) neurotransmitters, which can alter electrical synapse conductivity, as well as (ii) chemical activity which can influence nearby electrical coupling.

For example, Smith and Pereda [137] have early shown at mixed electrical and chemical synapses that chemical transmission can modulate the conductance of electrical synapses and thus regulate the degree of electrical coupling between neurons. Similar evidence was reported in the hippocampus more than a decade earlier [147].

Acknowledging that the effects of neurotransmitter modulation on the conductance of gap junctions has been extensively documented, Haas et al. [136] reported, in a dedicated review, *increasing evidence that electrical conductance in the mammalian brain can be influenced by the ongoing activity of neural networks*, thanks to local interactions with nearby chemical synapses. Given the widespread of the molecules and connexins involved in the process, the review suggested that this activity-dependent regulatory mechanism is likely a property of electrical transmission at large. Similar findings were also mentioned by other reviews [101, 142].

Overall, the literature shows the importance of the interaction between electrical and chemical synapses and its necessity[101] for normal brain functioning, beyond developmental phases. While more work is needed, the existing early evidence suggesting that chemical activity influences electrical coupling plays in favour of our theory.

Finally, it is worth noting that the recent review by Nagy et al. [138] has briefly mentioned the possibility of a backpropagating action potential, happening in a mixed synapse, and travelling backwards and laterally through nearby electrically coupled pre-synaptic neurons, “perhaps even promoting synchronization” therein, as partly shown by prior work of the same author[148, 149]. It turns out that gap junctions are crucial for a form of synchronized activity spanning populations of networks, and crucial for memory.

#### 3.4.3 Role in promoting network oscillations related to memory

We next present curious observations that press the need for further investigations. Central to our hypothesis is the proposition that backpropagated currents flow in reverse through coupled gap junctions, retracing the paths previously used during encoding, thereby activating distributed neural networks within the brain. If this phenomenon is as ubiquitous as our theory claims it to be—spanning cue-based attention, memory encoding, retrieval, to future episodic thinking— then its signatures should be prevalent and readily observable, e.g. via EEG or Dynamic Causal Modelling. One tempting candidate that satisfies all the above conditions is indeed the prevalent network oscillations[150, 151] involved in top-down processes related to memory and attention, and which synchronize distributed populations of neurons. And it is intriguing to note that those oscillations central to memory recall are disrupted when gap junctions are compromised. We review this aspect in the following.

Network oscillations represent synchronized neural activities across distributed regions of the brain[152, 150]. Many of these patterns, characterized by their frequency, amplitude, and phase, emerge transiently in response to specific cognitive or perceptual demands[151] such as attention and recall, or states (e.g. awake vs. sleep). Their amplitude measures the degree of neural synchrony, with a higher power indicating a more synchronized ensemble. Closer to our problem, gamma and theta rhythms, and their coupling [153, 154, 155], have been consistently associated in the literature with the complex processes underlying explicit memory formation, consolidation, and retrieval [152, 150, 156, 157, 153, 154, 158]. ^11^ Serving as integrative mechanisms, it has been suggested that such oscillations cause the reinstatement of memory traces stored in the cortex [156, 160]. By synchronizing neural populations, they would allow to (i) bind their information, for example to retrieve the contextual elements related to the cue (e.g. the temporal and spatial details of encountering an apple), as well as binding together various perceptual features [161] (e.g. an apple’s color and shape stored in distinct areas) [157, 156]. We successfully simulate a similar binding function in an artificial neural network in Sec. 5 where we reconstruct an image in a backward-propagation fashion from a small set of neural activations. Finally, theta oscillations and theta-gamma coupling in particular are believed to additionally (ii) structure memory retrieval of events in the correct order [156, 155, 162], an aspect that we leave for future work (cf. discussion).

If such oscillations are modulated by backpropagation-based top-down recall, and if gap junctions are the medium of backpropagated currents as we suggested earlier, then disrupting them should also disrupt oscillations. It turns out that abundant empirical evidence backs up this statement: in most of the documented cases above, gap junctions have been unequivocally implicated in generating and maintaining network oscillations. Specifically, it has been shown that whenever gap junctions were blocked or genetically removed, the synchronous oscillatory activity was disrupted or desynchronized [163, 164, 165, 166, 102, 102, 128, 129, 131].^12^.

In particular, focusing on our regions and functions of interest, a comprehensive literature review [167] confirmed the role of gap junctions (GJs) in general, and Cx36 to a large extent, in modulating network oscillations within the hippocampal formation, both in vitro and and in vivo. Interesting for future work, the effects, which were visible for both theta and gamma, varied depending on the experimental methods used to block gap junctions: Using pharmacological blockers such as carbenoxolone and octanol often led to simply suppressing oscillations all together, while gene knockouts that focus on electrical synapses, i.e Cx36 null mice, led to disruptions such as a significant decrease in power and frequency. Given that pharmacological blockers affect not only electrical synapses but also block other gap junctions, such as glial ones, this hints again to their possible role in the process. This aspect deserves careful future attention, especially in light of emerging reports about the role of astrocytes in gamma oscillations and recognition memory [97, 135].

In summary, the ubiquitous role of gap junctions and electrical synapses in fostering network oscillations [168] — which are essential for cue-based attention and memory across distributed brain networks — offers an intriguing avenue for exploring our hypothesis.

#### 3.4.4 Gap junctions: a crucial role in memory

If gap junctions play a role in the oscillations and synchronization that is necessary to form the distributed memories, then blocking them should also impair memory functions. Experiments with gap junction blockers and Cx36 null mice revealed severe memory impairments in such mice, in addition to other functional deficits including visual and motor [169, 166, 102, 170, 171, 172, 173, 174]. We here overview some examples of documented gap-junction-related memory impairments.

Frisch et al. [173] used Cx36 null mice to explore the role of gap junctions in memory and learning. Through behavioral tests like the Y-maze and object recognition, the researchers found that Cx36-deficient mice exhibited significant impairments in motor-coordination learning and memory, especially when faced with complex stimuli. For example, in the Y-maze test, these mice failed to habituate to a complex spatial environment even after 24 hours of retention. Interestingly, in object recognition tasks, the Cx36-deficient mice were unable to recognize objects after short delays of just 15 and 45 minutes.

Similar findings were reported in mice lacking Cx31.1[175], another connexin that is more and more suspected to be expressed in neurons in the brain. The authors have found that object recall was considerably impaired, after all delays, in addition to other altered memory-related behaviors, such as spatial novelty-induced exploration. Interestingly and worth future investigation, in relation to our hypothesized role of ACh as a trigger for the process (see Sec. 3.5 below), the authors found increasing levels of Acetylcholine Esterase release. Acetylcholine Esterase is indeed an enzyme that can break down ACh, ending the effect of cholinergic modulation.

Interestingly, but needing further investigation, Bissiere et al. [176] have found that gap junction blockers affected only (episodic) context-dependent memory (e.g. context fear memory) and not simple associative memories. Somewhat similar, Allen et al. [177] have shown that Cx36 lacking mice specifically exhibited deficits in short-term spatial memory when tested in a T-maze task. In this task, the mice were required to remember and alternate between left and right arms, a process that tests their ability to recall recent spatial information. On the other hand, their long-term spatial reference memory, which involves repeated training over several days to locate a submerged platform in a water maze, remained intact. Another study[178] on rats highlighted the role of gap junctions in the hippocampal CA1 area in the process of memory retrieval in state dependent learning. There the blockade of these channels by carbenoxolone disrupted morphine-induced memory retrieval.

Dere and Zlomuzica [179] reviewed the role of gap junctions in the brain, concluding that we need to revise the classic chemical-centered view of memory, to include the central role of gap junctions in episodic memory among others. Also deserving future attention, it is worth reminding that, beyond electrical synapses, other gap junctions such as those present in glial cells seem to play a crucial role in memory[180].

#### 3.4.5 Takeaway

As Nagy et al. [100] points out, the functional roles of electrical synapses “are still relatively few compared with the large numbers of reports on the pervasiveness of these synapses in various parts of mammalian CNS”. This limited understanding possibly explains, according to Nagy et al. [100], why the broader neuroscience community has not fully recognized the potential impact of these synapses on shaping neuronal activity. Our backpropagation-based recollection hypothesis offers a solution to this knowledge gap, suggesting a role for these synapses in crucial and ubiquitous functions ranging from cue-based attention to episodic memory.

Overall, all the references above confirmed the wide presence of electrical synapses and gap junctions more generally in various areas of the brain, including widely in the hippocampus and cerebral cortex, and with disruptions that are consistent with a possible role of gap junctions as the medium of backpropagated action potentials starting in the hippocampal formation and synchronizing large populations of neurons, causing reactivation or retrieval of information from the neocortex that stores them and that fired first when encoding them. Indeed, studies with gap junction blockers or genetical knock out confirmed the role of gap junctions in (i) synchronizing distributed large populations of neurons and (ii) explicit retrieval tasks, especially those that are short-term and context-dependent, intriguingly inline with our hypothesis.

### 3.5 Role of (cholinergic) neuromodulation as a trigger

As explained earlier, backpropagation-based recollection needs a trigger that could act as follows. Whenever the storage or recollection are deemed necessary, a neuromodulator is released in a diffuse non-selective manner to the areas where the traces are stored, from source layers to all earlier cortical areas where the traces are stored. Such neuromodulator would act in an activity-dependent manner, triggering the backpropagation-recollection from the sparse neurons that are firing at that moment. During a first encounter, this would favour cue-based attention: recalling the details of a particular cue and ignoring the rest. This would also help encoding as neuromodulation would favour the “rehearsal” of the backward paths while they are still “warm”, contributing to the coupling of neurons, to ease the later retrieval. At the time of retrieval, their role would be only to swiftly favour backpropagation, as briefly as the recall of the trace is needed. While more work is needed, a plethora of studies curiously corroborate the possibility that cholinergic modulation (through ACh), which we showed earlier to trigger backpropagated APs, could be such a neuromodulator.

#### 3.5.1 Triggering backpropagated APs

First, we remind as mentioned earlier in Sec. 3.1 that many studies have observed, in hippocampal pyramidal neurons, that cholinergic agonists enhance the backpropagation progressively, leading to stronger and stronger effect on subsequent APs [25, 26, 27, 28]. We focus in this section on subsequent effects of ACh that are inline with our framework.

#### 3.5.2 Agnostic diffuse cholinergic innervation of the cortex

Cholinergic modulation initiates in the basal forebrain from which cholinergic neurons innervate various areas of the cortex. ACh acts through both nicotinic (ionotropic) and muscarinic (metabotropic) receptors. Activation of these receptors can influence membrane potentials, affecting the firing patterns and synchrony between neurons. Huppé-Gourgues et al. [181] observed that the cholinergic projections from the basal forebrain to the primary and secondary visual cortical areas (V1 and V2) were, interestingly, not organized retinotopically suggesting that activation of the basal forebrain does not selectively stimulate specific areas of the cortex. Instead, they result, *each time*, in a broader or more diffuse activation, hence influencing a larger portion of the visual field rather than a specific region. This is consistent with our hypothesis that such neuromodulation is diffuse and its effects activity-dependent.

#### 3.5.3 Triggering network oscillations related to memory

Second, if ACh serves as the trigger for backpropagating action potentials, which in turn triggers wider synchronized network activities related to memory, then inhibiting its function should also alter both network oscillations and explicit memory formation and recall. We find again a plethora of studies that curiously back up these statements too. Indeed, the medial septum and diagonal band of Broca complex innervate also the hippocampus, and this major cholinergic input was shown, already a while ago, to induct network oscillations that are similar in frequency to those important for memory functions [29, 182]. Indeed, early work showed correlations between the release of ACh and increase in oscillatory power in vivo [182]. The knockout of specific ACh receptor subtypes further provided early evidence for this scenario [29].

Later work provided more evidence and more details. For example, Betterton et al. [183] used Carabachol (a muscarinic ACh receptor agonist) as a proxy to show that ACh modulated the power of existing gamma oscillations in the hippocampus through muscarinic M1 receptors, without affecting their frequency. This modulation interestingly works in “‘both directions” depeding on the cholinergic dose: low ACh leads to an increase, but a too high ACh conversely leads to a decrease of oscillation power. This is similar and complementary to the conclusion reached earlier by Wang et al. [30], but using nicotinic receptors. The study showed the modulation of gamma oscillations in the rat hippocampal CA3 area by nicotine (a nicotinic acetylcholine receptor (nAChR) agonist). Similarly, Wang et al. [30] have found that low concentrations of nicotine enhanced gamma oscillations, while high concentrations reduced them.

Similar results were obtained on anesthetized mice using optogenetic stimulation of septal cholinergic neurons [184]. There, the stimulation was shown to suppress sharp wave ripples and enhance both gamma, and theta oscillations in the hippocampus. Little effect was found on the awake mice, when the background had already high theta power. Similarly, Dannenberg et al. [185] utilized optogenetic stimulation on mice to demonstrate that cholinergic neurons in the medial septum and diagonal band are crucial for generating and pacing theta-band oscillations in the hippocampal formation. Interestingly, this neuronal activity was specifically observed during exploration, novelty detection, and memory encoding processes.

Beyond triggering oscillations that, as we showed earlier, are key for memory and attention, recent work has more directly shown that the oscillations triggering process coincides with key cognitive functions such as cue detection. As we will argue later, cue detection involves a form of retrieval, and Howe et al. [186] showed that ACh release in the prefrontal cortex promotes gamma oscillations and theta-gamma coupling during such cue detection.

Overall, the literature confirms the selective role of cholinergic signaling in the modulation of network oscillations that are known to be crucial for cognitive processes such as memory and attention.

#### 3.5.4 Triggering memory encoding and retrieval

While earlier work popularized the idea that cholinergic manipulation had less impact on memory retrieval compared to encoding[187, 188, 189, 190], more and more recent evidence suggests a clear role in the retrieval of (both recently and remotely acquired) memory traces [191, 192, 193, 194, 195, 196].

As an example of early work, Miranda and Bermúdez-Rattoni [187] used tetrodotoxin (TTX) to disrupt the activity of the nucleus basalis magnocellularis (NBM), a key region for cholinergic modulation, showing that this disruption impaired the formation of new taste aversion memories in rats. The authors then found no disruptive effect on memory retrieval, once memories were already formed. Along the same line, Rogers and Kesner [189] concluded, after studying the effects of scopolamine (a cholinergic antagonist) and physostigmine (an acetylcholin esterase inhibitor) on fear conditioning in rats, that different levels of ACh had different impacts on encoding and retrieval: low levels of ACh impaired encoding but not recall. Very high levels of ACh even impaired retrieval, the study have found. Hasselmo and McGaughy [190] also stressed the role of ACh mostly in encoding suggesting that high levels of ACh enhanced attention to sensory input, facilitating the encoding of new information, while low levels of ACh, on the other hand facilitated memory consolidation.

Since then, numerous studies revised the role of ACh to include also retrieval. Soma et al. [195] focused on the role of ACh in the retrieval of well-trained memories formed through repetition and daily routines (as opposed to back-then frequent studies that focused onl on recently acquired memories). The authors showed that the administration of scopolamine resulted in the impairment of task-related associative memory in rats, and underscored the essential role of muscarinic acetylcholine receptors (mAChRs), activated at normal acetylcholine levels, in the retrieval of well-trained memories.

Using both pharmacological and genetic methods, Leaderbrand et al. [191] further challenged the older view that ACh predominantly supported learning, revealing its crucial role in retrieval as well. The research demonstrated that both encoding and retrieval of contextual memory required the activity of mAChRs in the dorsal hippocampus (DH) and retrosplenial cortex (RSC), and further elucidated distinct roles of mAChR receptor subtypes in this complex process: For memory formation, both DH and RSC primarily utilize M1/M3 receptors. However, for memory retrieval, while DH continues to rely on M1/M3 receptors, RSC shows a notable shift, employing M2/M4 receptors for recent memory retrieval and all receptor subtypes for remote memories.

Moosavi et al. [194] also showed that scopolamine impairs memory retrieval in a passive avoidance task (animals learn to avoid an environment where they previously received an unpleasant stimulus). The study found in addition that scopolamine induced changes in molecular markers in the hippocampus such as MMP2 and MMP-9 (Matrix Metalloproteinases 2 and 9), which are enzymes linked to hippocampus-dependent memory and neurological disorders like Alzheimer’s Disease.

Sun et al. [192] further demonstrated that scopolamine administration in mice significantly impairs the retrieval of spatial information in the hippocampus. This impairment was notably reflected in the reduced accuracy and stability of spatial representations by hippocampal neural ensembles. The study also observed a decrease in the number of active neurons, their firing rates, and the quantity of place cells. Unlike previous work which measured the retrieval performance using behavioral metrics, this paper analyzed the impact of scopolamine on the functional integrity of place cells (which are typically integral to forming a stable and accurate environmental map). Indeed, following scopolamine treatment, there was a notable disruption in this mapping accuracy. Using a Bayesian decoder to estimate the mice’s location based on place cell activity, the researchers observed a substantial increase in decoding errors after the administration of scopolamine.

Olson et al. [193] recently provided further evidence, and corroborated the findings of Moosavi et al. [194]. Through scopolamine administration, Olson et al. [193] showed not only disruption of spatial memory retrieval but also other associated biochemical changes in the hippocampus (minimizing retrieval-induced alterations in MMP-9, complementing the work of Moosavi et al. [194]).

Last, noticing how the early ideas that high levels of ACh impaired hippocampus-mediated retrieval[189] conflicted with the fact that high levels of cortical activity were necessary[197] for attention-demanding recollection, Decker and Duncan [196] suggested in a recent review that rapid cholinergic mechanisms causing swift neural oscillations may resolve these apparently conflicting retrieval demands. This is inline with our hypothesized role as a transient trigger. We next show evidence that such transient signals are also key to cue-based attention.

#### 3.5.5 Triggering cue-based attention (retrieval)

Cue detection is a critical cognitive process that requires selective attention to specific external cues to guide behavior. This attentional mechanism can be seen as a form of immediate cue-based recall, promoting the activity of selected neural networks uniquely signaling the cue, thereby focusing attention on it and reducing responsiveness to other background signals. It turns out that phasic cholinergic release or cholinergic transients, which are brief, discrete, release events of ACh, have been closely associated with the detection of cues in attention tasks, suggesting a possible role as a trigger inline with our hypothesis.

Howe et al. [198] provided converging evidence, from rats and humans, that cholinergic transients trigger the shift from the state of passive monitoring of cues to an active cognitive state of cue-directed response. The authors found in particular, in rats, that ACh release in the Preforntal Cortex happened in trials where the signals were correctly identified (while it didn’t happen in case of miss or during consecutive correct identifications). Howe et al. [186] later employed a comprehensive approach combining subsecond measures of cholinergic signaling, neurophysiological recordings, and pharmacological blockade of cholinergic receptors to elucidate the role of ACh in cue detection within the prefrontal cortex. Building upon prior work [198], the study demonstrated that phasic transient increases in ACh release are critical for effective cue detection. Specifically, these increases in ACh release were temporally aligned with enhanced neuronal synchrony across various frequency bands and the emergence of theta–gamma coupling. Notably, the study found that both nicotinic and muscarinic receptors play distinct but complementary roles in modulating these neural oscillations during cue detection.

Sarter et al. [199] employed amperometric recording techniques to measure choline-sensitive transients in response to attention tasks, revealing a direct association between brief, phasic acetylcholine release events and the detection of cues. Their findings indicate that these cholinergic transients are crucial for cue detection, independent of reward outcomes, and play a significant role in shifting attention from monitoring to active cue detection.

Sarter et al. [199] also thoroughly reviewed the role of phasic cholinergic transients in rodents concluding that they are essential for the transition from monitoring to cue detection. Interestingly for our activity-dependent claim, they found that more salient signals are more likely to trigger cholinergic transients. The review concluded with a set of open questions such as: (i) what cognitive state triggers the occurence of the transients, (ii) what is the precise temporal dynamics of the transients, specifically the timing relative to the signal cue and glutamatergic input, and finally (iii) the interaction between longer-term cholinergic neuromodulation and the generation of transients. *Under our framework, neurons representing the most salient features fire more strongly. Then cholinergic neuromodulation favours backpropagated APs proportionally to such firing rates. The resulting currents synchronize populations of neurons related to the salient cue, leading to an increase of activity of those neurons representing the cue to pay attention too, and a relatively reduced activity of the rest of the stimuli*.

#### 3.5.6 Takeaway

In addition to the references above, Haam and Yakel [31] thoroughly reviewed the role of cholinergic modulation in memory, providing a comprehensive synthesis. First, it confirms that cholinergic modulation paces the firing activity of hippocampal neurons and modulates the oscillations therein. Second, it concluded that ACh’s effect is specific to hippocampus-dependent declarative memory only and that procedural procedural memory is not impacted. For example, Flesher et al. [200] have shown that ACh release in the medial prefrontal cortex and hippocampus is significantly higher during trace conditioning tasks (a declarative memory) compared to delay conditioning. Third, and interestingly, the review spotted seemingly contradicting effects of ACh on the hippocampus, with many works showing that ACh release in CA1 potentiated the pathway between the CA1 and CA3 areas of the hippocampus, and many others showing instead an inhibitory effect. The authors hypothesized that the effect of ACh could be timing-dependent. While further investigations are needed, this is inline with our backpropagation-based recollection, where the effect of neuromodulation release on backpropagation is activity-dependent. Last, the review reported studies showing an atrophy in the cholinergic system of early Alzeihmer Disaese (AD) patients, as well as others showing changes in cholinergic receptors.

Overall, the literature we reviewed agrees on the crucial role of cholinergic modulation during the first encounters, for both memory encoding and cue-based attention, which as we explained earlier is another form of immediate recall under our our framework. If our hypothesis is correct, this allows to rehearse the backward pathways and strengthen cell coupling while the pathways are still “hot”. Next, while earlier work has shown a role of ACh mostly during first encounters, more and more work supports the role of rapid cholinergic release also in attention-demanding retrieval tasks. This is also inline with the fact that cholinergic release is low in states like SWS and is high in wakefulness states and REM sleep (where dreams, a sort of cortical retrieval, happen most).

Finally, it is worth mentioning that it could be also possible that other neuromodulators play a role in triggering retrieval. In this paper, by focusing on ACh, we have shown that neuromodulation more generally (and cholinergic modulation in an intriguing fashion) can act in ways that make our backpropagation-based recollection theory biologically plausible.

### 3.6 Summary of arguments in favor

We briefly recapitulate, as simplified in Table. 1, arguments in favor of backpropagation-based recollection. Fig. 3 also summarizes the found evidence on the toy example of Fig. 2. First, as seen in Sec. 3.1, activity-dependent backpropagating APs are biologically plausible. Moreover, these APs are stronger when neurons are firing, a necessary condition for our hypothesis. Second, (cholinergic) neuromodulation can enhance such backpropagation in a selective and progressive manner, thus acting as a switch to inhibit it or to strengthen it. This “feature” is necessary as subjects can control whether or not to favor retrieval upon exposure to a cue, and as, obviously, not all cues lead to recall. Interestingly enough, this modulatory phenomenon on backpropagated-APs was observed in hippocampal cells, a crucial area for explicit memory retrieval [76, 80], especially known [74, 73, 80] to contain indexes that allow to retrieve memories stored in other cortical areas. We further found that sparse neurons, similar to our pointer neurons or indexes, have been also well-documented in these areas [8, 93]. Even more, a top-down reversal of information flows was observed therein (Sec.3.2.2) following the bottom-up perception of a cue. This brings us to our next assumption, which is that retrieval is the reactivation of the same neural ensembles used during encoding. There also, a large body of optogenetic and neuroimaging-based evidence confirms it. After all, all optogenetic experiments cited above tagged the specific neural ensembles that were active during encoding, to later observe them during retrieval.

**Fig 3.**
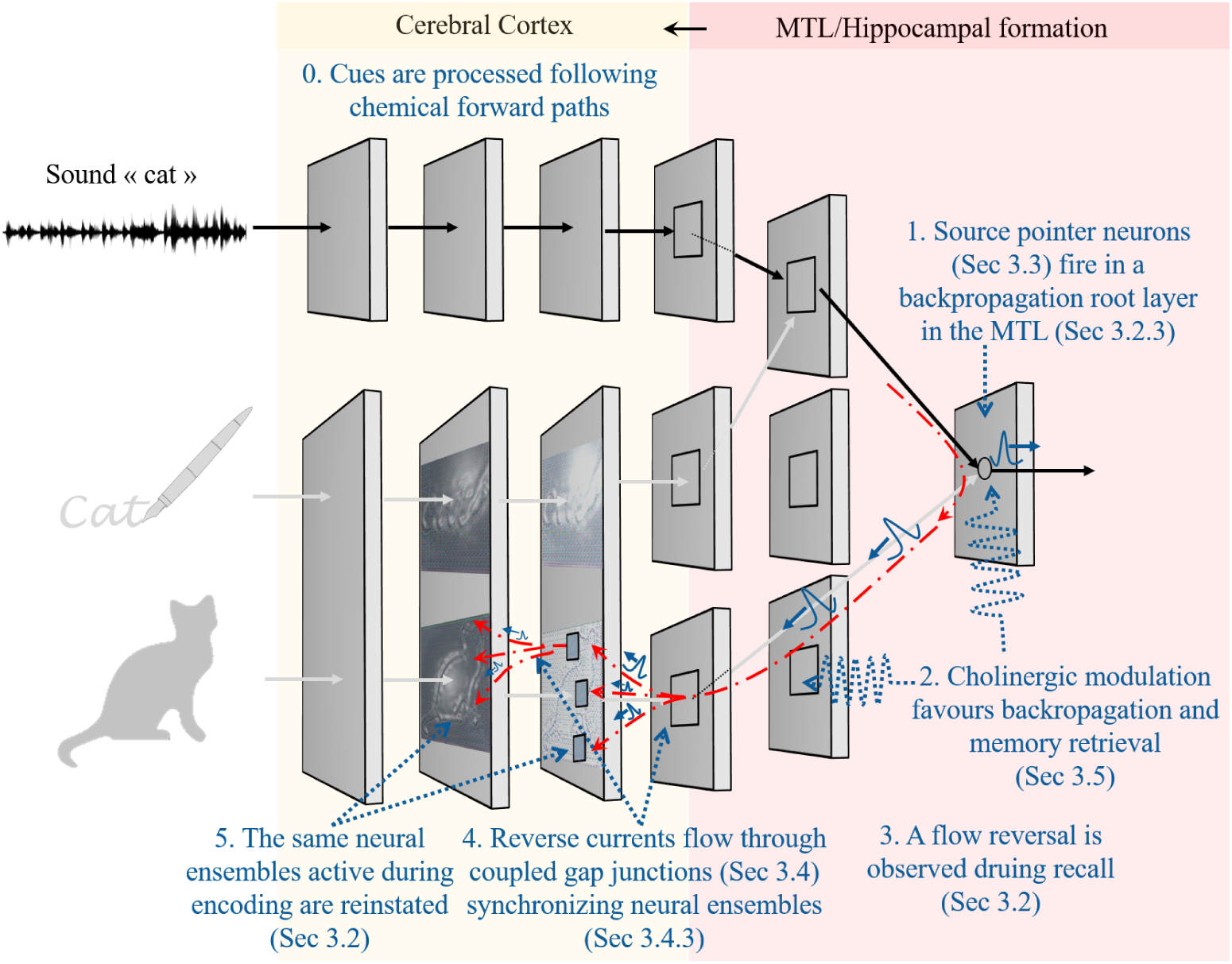
A biologically plausible *illustration* of the toy example of Fig. 2 during recall, under the light of evidence we found in the literature.

Next, we have found that backpropagated APs can have far-reaching effects, and that gap junctions which were found to be tremendously more prevalent in the cortex than previously thought, could be the medium through which backpropagated currents could travel, synchronizing large populations of neurons and favouring retrieval. Indeed, blocking gap junctions lead to memory impairments as well as disruptions of network oscillations. Finally, a plethora of evidence links ACh to our hypothesized role of trigger to the backpropagation-based recollection.

Finally, beyond cue-based recollection and attention, backpropagation-based recollection is a candidate mediator of a broad range of generative tasks such as mind-wandering, creative thinking and future episodic thinking whose neural correlates are still also open questions [201, 202, 203].

Next, we computationally show that backpropagated APs are as effective as machine learning in associating names to visual input. Later, we show that they can reconstruct an image from a set of sparse activations.

## 4 Efficiency in object classification

We now simulate our hypothesis, assessing whether it is a computationally efficient strategy for the task of retrieving object names using their image as a cue. For this, we leverage existing artificial Spiking Neural Networks (SNNs) trained with *Spike Timing Dependent Plasticity* (STDP) and simulate a “teacher” that simultaneously trains the SNN with images and their corresponding names. Then during test, backpropagated APs are used to retrieve the right name. We compare the accuracy of our hypothesis against a machine learning classifier.

### 4.1 Materials and methods

We first describe the recent models we build on, acknowledging the aspects in which they lack plausibility (Sec. 4.1.1). We then describe how we use them to model our hypothesis (Sec. 4.1.2). We finally detail our experiments and evaluation protocol (Sec. 4.1.3).

#### 4.1.1 Image classification with existing Spiking Neural Networks

SNNs are biologically-inspired computational models in which spiking neurons communicate through individual spikes that propagate from one neuron to the next. Such spikes simulate APs, happening when the neuron’s membrane potential crosses a certain threshold. The artificial SNNs we leverage in this paper simulate a simpler version of this process, yet still offering higher biological plausibility [204] compared to other artificial models. Indeed, for training, the SNNs we leverage use the more plausible STDP learning rule [205, 7, 206, 207, 208] where synaptic weights are updated according to the relative spike times of pre and post synaptic neurons: if the pre-synaptic spike occurs slightly before the post-synaptic spike, then a persistent strengthening of synapses called long-term potentiation (LTP) occurs [209]. In the other case, the result is a long-term depression (LTD).

Two recent models provided background for our simulations [3, 4]. We build on that of Kheradpisheh et al. [3] which achieves impressive accuracy on simple datasets. We reuse “as-is” its feature extraction layers illustrated in the “feature learning (STDP)” upper part of Fig. 4. The latter consists of consecutive neural processing layers: the first is a temporal coding layer that somewhat mimics retinal ganglion cells firing moments, followed by a cascade of convolutional and pooling layers to extract visual features. In details, the first layer encodes the input signal into discrete spike trains in the temporal domain, it uses Difference of Gaussian (DoG) filters.^13^ This layer detects positive and negative contrasts in the input image and encodes them in spike latencies, according to their strengths. Next, each neuron in the convolutional layer receives input spikes from the neurons located in a certain window and emits a spike when its potential reaches a specific threshold. Pooling layers perform a nonlinear max pooling operation in which they only propagate the first spike emitted. In this model, STDP learning only occurs in convolutional layers and is done layer by layer. For each image presented to the neural network, there is a “competition” between neurons of a convolutional layer and those which fire earlier trigger STDP and learn the input pattern. Finally, the last layer is a global pooling layer which performs a global max pooling. The role of feature extraction layers is just to learn visual features: they are trained without labels, by propagating many images through the layers and adjusting the weights with STDP.

**Fig 4.**
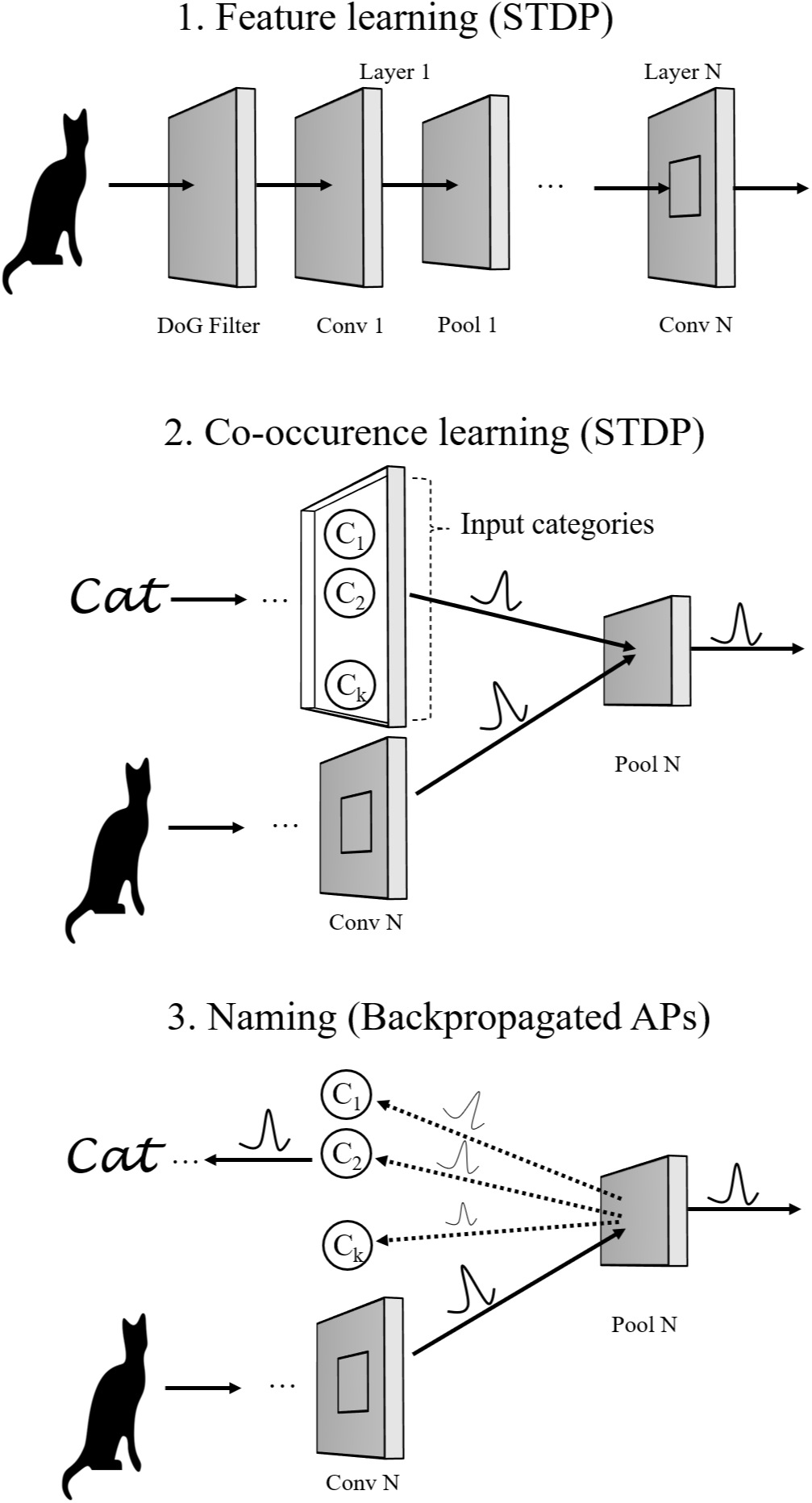
End-to-end learning, simulated with the proposed SDNN with its three main parts. Backpropagation-based recollection happens during the third step: naming. As exemplified in the figure, after showing the image of a cat, backpropagated APs travel back to the input categories layer. The C2 neuron got a high score, successfully signalling that the image corresponds to a cat.

Next, unlike our hypothesis, the output of this final layer is used in the model of Kheradpisheh et al. [3] to train a linear Support Vector Machine (SVM) classifier. The SVM classifier is of course not biologically plausible but the goal of Kheradpisheh *et al.* was only to assess the ability of SNNs/STDP to extract salient features that are good enough to discriminate images. And they actually found that they were good enough in terms of classification accuracy: their implementation reached 99%, and 98.4% of accuracy in the face/motorbike and MNIST datasets, respectively. We reproduced similar^14^ results on the face/motorbike dataset building on Perez’s available implementation [210].

##### Plausibility

Finally, before using Kheradpisheh’s model [3] as a basis, we briefly acknowledge its lack of plausibility, arguing though that it does not impact the simulation of our hypothesis. Despite being unsupervised, and despite using STDP, the model uses only a spike-time neural coding (i.e. rate is not modeled). It also uses convolutional neural networks with weight sharing, which is also biologically not plausible.^15^ These “problems”, however, do not impact our hypothesis since we are only interested in the last layers, where the APs backpropagation will actually initiate. All we need, is a set of decent learned features therein.

Since these models trained only with unsupervised STDP achieve good performance on what was 20 years ago a difficult problem, we assume that they extract visual features of fairly good quality. The latter are obviously not perfect, i.e. accuracy does not reach exactly 100% even on the simplest motorface dataset. Nonetheless, we set out to verify whether our hypothesis, together with STDP, can successfully use these same features to find the right name association. For fairness, we compare our performance results to those of the SVM classifier.

#### 4.1.2 Our three-step model

Under our hypothesis, successful name association comprises three steps, two for learning and one for recollection, as modeled in Fig. 4. The first *feature learning* step is completely unsupervised and learns through repeated exposure to visual stimuli to extract salient features (e.g. lines, shapes etc) to discriminate visual content. In the brain, such learning is supposed to happen early in life. And if, for some reason or another, one is not exposed enough to visual stimuli, such learning does not happen, which leads to cortical blindness. For this part, we reuse the SNN model described above [3].

The second step is a semi-supervised learning one, whereby a teacher shows a learner an image and the right name that refers to it. We call this *co-occurrence learning*. Humans for instance can learn from a single example to map a new object to its new name. When no external reinforcement happens, exposure may need to be repeated until it is remembered. We model this step by simply adding a new “categories layer” and emulating the right spike each time an image is propagated through the SNN, i.e. generate a spike for “cat” category neuron, while the cat image is propagated through the SNN.

The third step is simply the recollection of the name. It is in this step that an AP is backpropagated from *each neuron* in the backpropagation root layer, towards the categories layer. There, the neuron(s) which receive the highest “vote” signal the network decision. We now describe the three steps of Figure 4 in more details.

##### Feature Learning

This step uses almost as-is the SNN model above [3], the reader can hence refer to the original paper for more details. This phase starts with the input image being encoded into discrete spike events in the temporal domain. Each “spatial position” or pixel in the input image will have its own spiking time. Such time-encoding process is performed by using Difference of Gaussian (DoG) filters, where spike times are computed according to the DoG filter’s output. More precisely, let *r* be the value at a certain position index after having applied the DoG filter. Then, the firing time *t* is defined to be 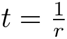. This corresponds to encoding higher contrast areas of the image to lower spike times (i.e., latency is inversely proportional to the contrast). As a result, each single image is transformed into several waves of spikes that propagate, one by one, through the layers: spikes that signal higher contrast areas being the first to enter the network. Next, Convolutional layers are arranged in a feedforward manner. Between two consecutive convolutional layers, a pooling layer performs a max operation to compress visual data and provide translation invariance. The task of a neuron in a pooling layer simply consists in propagating the first spike received from a receptive window of the previous convolutional layer. Neurons in all the convolutional layers are *non-leaky integrate and fire* neurons. They integrate input spikes and emit a spike as soon as they reach their threshold. The latter is a hyperparameter to set. Immediately after a spike occurs, weights are updated accordingly, using a simplified version of the STDP learning rule. Let *i, j* be the indices of the post and pre-synaptic neurons, respectively, and let *t_i_, t_j_* be their corresponding spike times. The synaptic weight *w_ij_* is updated by adding a modification factor Δ*_ij_* computed as follows according to a simplified version of STDP [3, 212].

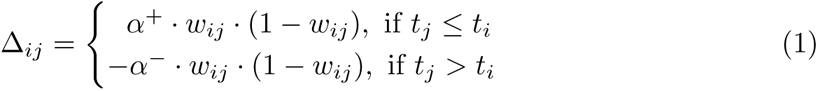

*α*^+^*, α^−^ ∈* ℝ*_≥_*_0_ are two parameters that specify the learning rate or by how much the weights are changed. The latter factor impacts a lot the learning. Indeed, small values would lead to a slow learning process, they simulate a neural network that is confident in its prior “beliefs and decisions” (weights). High values allow to learn very fast information about the current stimuli, but they can have as a consequence to “forget” what they learned with previous stimuli. Note that this simplified version of the STDP rule does not take into account the absolute time difference between post and pre-synaptic spikes. What matters instead is only the order, or the sign of the difference. In practice, this is not a problem for our model.

This feature learning process goes on, by propagating training images one by one. Each time the different spiking waves of a new image have been fully processed, and weights updated and stored, the potential of each neuron is reset to 0, preparing for the next image. Initially, the synaptic weights are chosen at random from a normal distribution with some mean and standard deviation^16^. The STDP rule ensures that they always remain in the range [0, 1]. Within each image, learning is done layer by layer: learning at layer *ℓ* begins when learning at layer *ℓ −* 1 has terminated.

The intent of this feature learning phase is to learn the synaptic weights of each neuron in all the convolutional layers. As observed previously with this SNN model [3], neurons in the first layer converged to the simple four oriented edges. Neurons in the successive layers learned more complex ones by integrating spikes from previous layers. We remind that this phase is totally unsupervised as the network only learns frequent features associated with images and it requires no knowledge about the input image categories. The next learning step includes such categories, we qualify it as semi-supervised.

##### Co-occurrence learning

Once the SNN has learned the right weights and hence visual features, the second co-occurrence learning step can begin. For this step, we assume as per our hypothesis, that there is a layer which encodes the object categories or names (equivalent of sparse neurons that respond to image or sound signifiers or both). The latter layer is connected, as shown in the figure, to the last layer of the image processing network. Then, an image (e.g. a cat as illustrated in the figure) is propagated through the image processing neural network, while at the same time, the neuron^17^ which represents its signifier or name is activated simultaneously.

In more details, as in the previous phase, train images are considered one by one. Using the weights learned in the first phase, each image passes through the network until it reaches the last layer where a max pooling operation is performed. During co-occurrence learning, a neuron of the last pooling layer would thus receive two spikes: one propagated by the neuron associated to the class of the image, and one input from the last convolutional layer. This entire process simulates the teacher that simultaneously shows the image and its name.

Note that, implementation-wise [213], this is equivalent to having a matrix with the same shape as the last pooling layer that is associated to each image category (one weight per category neuron per neuron in the backpropagation layer). As in the first phase, weights are initially random and are updated only using the STDP learning rule, as defined in (1). As a result of this STDP learning, according to the order of post and pre-synaptic spike times and to the index of the spiking neuron, what will happen is the following: the weight matrix of the correct image category is always strengthened (LTP), while the weight matrices of the other categories will be weakened (LTD).

Contrarily to the previous one, this phase is supervised in that it requires knowledge of the image category in order to be able to link it with the corresponding image features. Indeed, the aim here is to learn associations between the previously learned features and image categories by using only the simple STDP rule. At the end of the second phase, training has completed and the SNN can proceed to the naming task using the backpropagation principle.

One extreme version of this second phase is what is called one-shot learning: the network is given only a single example of each category. We will vary the number of such training examples in Sec. 5.2, effectively trying one-shot and few-shot learning scenarios, hence why we consider this task to be “semi-supervised”.

##### Naming

Once the *learning* is done with the previous two phases, we are ready now for the *naming* task, following the principles of backpropagated APs.

In this task, the image to name is first propagated through the fully trained neural network until spikes start to happen in the last pooling layer, which is our backpropagation root layer. We consider that all neurons in this layer are source pointer neurons. This means, as per our hypothesis, that we allow them (whenever they fire) to send backpropagated APs, modulated by the pre-synaptic weights learned in the previous second step, to the previous layer that encodes the labels or names. Neurons in the “categories/signifiers” layer will integrate such received signals and the category which has the highest vote is the retained name for the image. Namely, let *C_i_* for *i* = 1*, .., k* be the class associated to the neuron with the highest class score. Then, *i* is chosen to be the class the image belongs to. It is this score that can be used as the accumulated potential that brings the right neuron closer to its firing threshold, leading when it fires to the class decision^18^.

We show later in the next section that such a simple mechanism allows to label the images as accurately as the SVM. More interestingly, by using high learning rates during the first co-occurrence (e.g. neuromodulating to increase synaptic strength), it is possible to learn to name objects, with maximum accuracy, by showing the neural network only a single instance of the image class; an extreme learning task in which the SVM classifier seems to have more difficulty.

#### 4.1.3 Experiments and evaluation protocol

We design experiments to evaluate the computational efficiency of our three-step SNN model in associating a name to an image. As a baseline, we consider the performance of the SVM machine learning classifier on top of the learned visual features, exactly as was done by Kheradpisheh et al. [3]. We first present our evaluation tasks and training protocol, before describing how we hyper-parameterize our models.

##### Evaluation tasks and training procedures

We evaluate the efficiency of our hypothesis on two tasks of increasing difficulty: *(i)* classic learning from many examples, and *(ii)* one-shot learning after seeing only one example from each class. Similarly to Kheradpisheh et al. [3], we use the Caltech motor face dataset [214] considering two classes: Faces and Motorbikes. The dataset as employed by Kheradpisheh et al. [3] comes with a single-split of the 398 images from each class into 200 for training and 198 for testing. For both tasks, we consistently use the set of 200 training images for our first feature learning step.

For task *(i)*, in addition to the *single split* used by Kheradpisheh et al. [3], we further perform around 50 new random splits, always keeping the same proportions (i.e. 200 images for training and the rest for testing).

In task *(ii)*, we use only one image from each class for training. To get representative results, we repeat this experiment 1500 times, picking each time a different pair of (motor, face) image from the training set. We perform the test on the entire test set of 198 images.

In both tasks, in addition to verifying whether our hypothesis “works at all”, we compare its accuracy to that of Kheradpisheh et al. [3], i.e. the same feature learning as step 1 above, followed by an SVM classifier on the feature vectors.

Finally, we perform our experiments on a server with 5 Intel 2.10 GHz CPU, 32 GB of memory and a GPU Nvidia Tesla P100 SXM2 with 16 GB of dedicated memory.

##### Model calibration

We are not aware of an efficient biologically plausible method to set the hyperparameters of our spiking neural network models (learning rates, spiking thresholds etc.). As this is still an open research question, similarly to Kheradpisheh et al. [3], we programmatically search for best parameters.

For step 1, feature learning, we perform a grid search to find the parameters [213] that maximize the performance of the SVM classifier on the training set. This allows us to both reproduce the performance results of Kheradpisheh et al. [3] and ensure that the learned features are of good quality. We converge to *α*^+^ and *α^−^*of 0.007 and 0.003 respectively, for the feature learning layers. The firing thresholds for the first, second, and third convolutional layers are set to 6, 21, and 10. Similar to prior work [3], max pooling is not performed for this Caltech dataset because images have low resolutions. Our last layer uses thus a pooling window of size 1×1. Finally, the synaptic weights of the class matrices are initialized at random from a normal distribution with mean 0.59 and standard deviation 0.05.

For step 2, co-occurence learning, we perform another search, always using the training set, to set the STDP learning rates of the last co-coccurence based learning layer. For the one-shot learning task, since we only have a single example, we cannot afford to increase the learning rate in small increments: there is no repeated exposure to images and their names that will slowly adjust the synaptic weights. Instead, much higher learning rates are needed to properly set the weights correctly from the first tryout. Note that suddenly increasing the learning rate, this way, is biologically plausible: it can be seen as the result of a neuromodulatory action that strengthens a connection so as to “memorize” quickly, without the need for repeated exposure. To understand the impact of such modulatory effect and calibrate the STDP learning rates, we start from *α*^+^ = 0.007*, α^−^* = 0.003 as a basis and multiply them by some factor *λ*. We then vary the number of images shown during training and study the impact of scaling up and down he learning rates on the accuracy.

More detailed results will follow (Fig. 7), but as intended, we find that considerably increasing the learning rates allows to achieve high accuracy in the one-shot extreme task. For instance, *λ* = 65 yields good results among many other similar values, so we pick it to set the STDP learning rates for this second task, i.e. *α*^+^ = 65 *×* 0.007*, α^−^* = 65 *×* 0.003. On the contrary, for the first task, learning from many examples, we found that scaling down both *α*^+^ and *α^−^* yielded slight improvements: we accordingly set them to 0.00399 and 0.00171 respectively (*λ* = 0.57 but other values were suitable too.).

Not the topic of this paper, but these findings hint to a possible successful learning strategy in which the learning rates are decreased over time, starting with high rates during the first encounters (i.e. novelty-based modulation to learn quickly) followed by decays, to consolidate the learning and preserve the learned weights from future bad examples.

## Code Accessibility

The datasets and code for this study can be found in the following git repository: https://github.com/bendiogene/recollection_hypothesis.

### 4.2 Results

We now overview our results.

#### 4.2.1 Backpropagated-APs allow to separate classes

First, more qualitatively, Fig. 5 shows the scores obtained by each class neuron for both train and test datasets. Recall that these scores correspond to the “votes” received through backpropagated APs from the last layer (during inference, after the end of training steps 1 and 2). Each point in the figure corresponds to an image, with motorbike points placed on the left and faces on the right. Values in the ordinate represent the difference between the scores associated to the Motorbike and Face neuron classes. Therefore, images with positive values are associated to the Motorbike class, while images with negative values are associated to the Face one.

**Fig 5.**
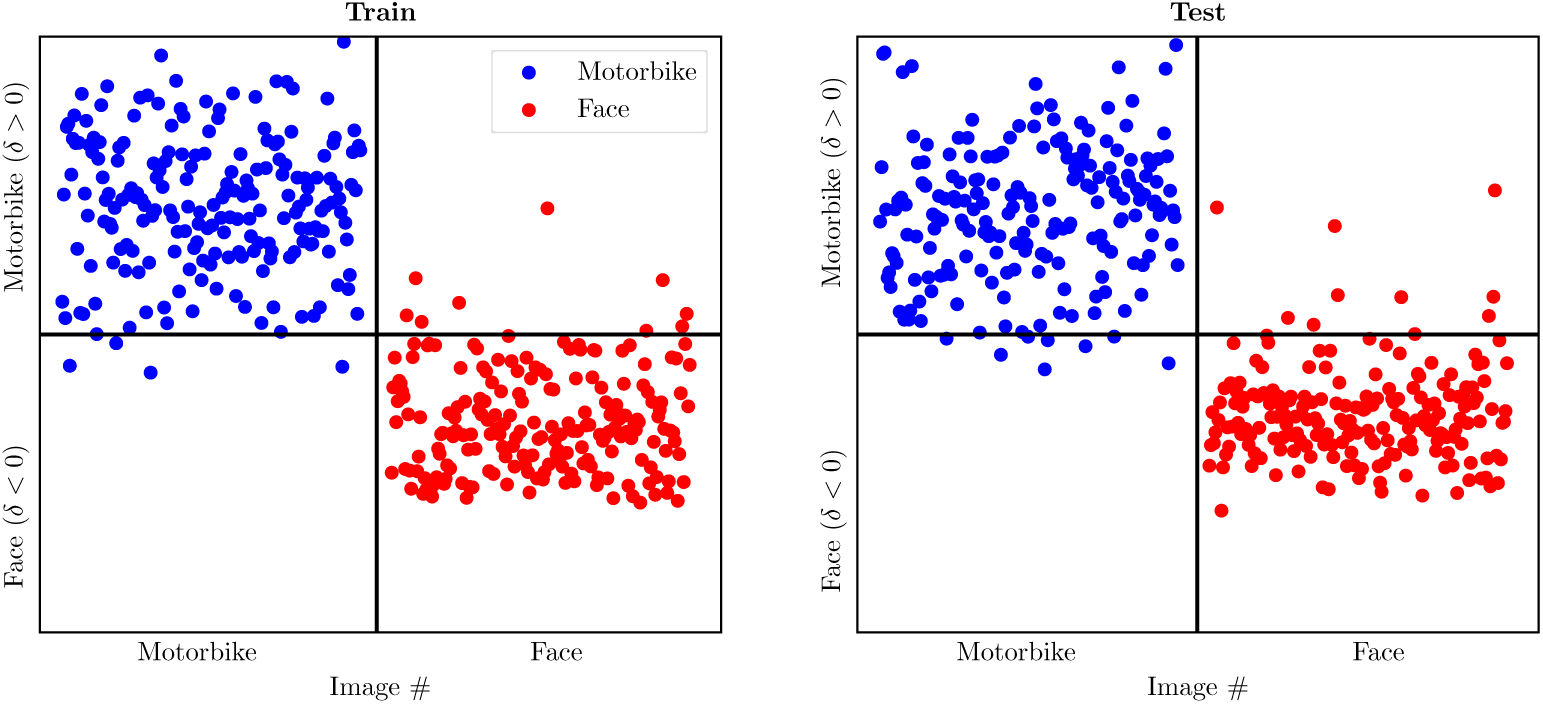
Illustration of computational efficiency of backpropagated APs. Difference between the class scores for each image in the Train (Left) and Test (Right) datasets. The class scores allow for separation.

As it can be observed, backpropagated APs allow to separate the two classes for most of the images. This already shows the computational efficiency of our hypothesis. For some of the images however, this distinction is not clear. As we discussed earlier, this could be due to the SNN feature extractor which, although performing, does not yet learn good representations. To shed more light on this, we next use the SVM as a baseline.

#### 4.2.2 Task 1: backpropagated-APs compete with the SVM classifier

Fig. 6 opposes the accuracies (on test set) of backpropagated APs and those of the SVM classifier, accross the 49 different train/test splits. The figure shows a Gardner-Altman paired estimation plot, using a Tufte slopegraph linking the pairs of accuracies obtained on the same split (instead of the classic swarmplot) [215]. Overall, using the backpropagation-based recollection, we reach an average accuracy of 96.8% less than 1% behind the SVM classifier, 97.6%. This means that backpropagated APs can not only capture some useful signal, but their accuracy is competitive with a supervised machine learning classifier such as SVM. Further, as shown by slope graphs, in a non negligible number of train/test splits, the backpropagation-based recollection yields higher accuracy. With future enhancements to the modeling of STDP learning, these results will likely improve.

**Fig 6.**
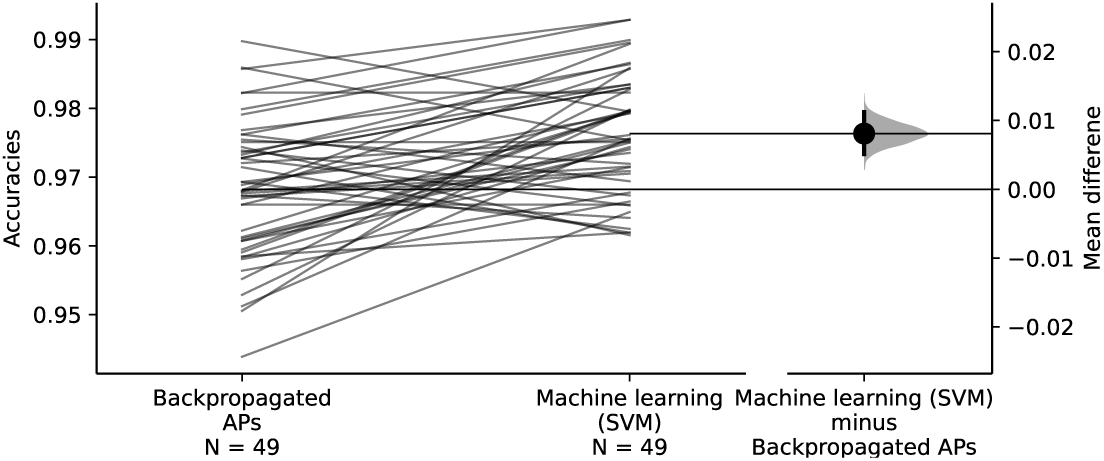
Backpropagated APs compete with the SVM classifier. Gardner Altman paired estimation plot, opposing the accuracies of backpropagated APs vs. SVM when trained on 200 images.

**Fig 7.**
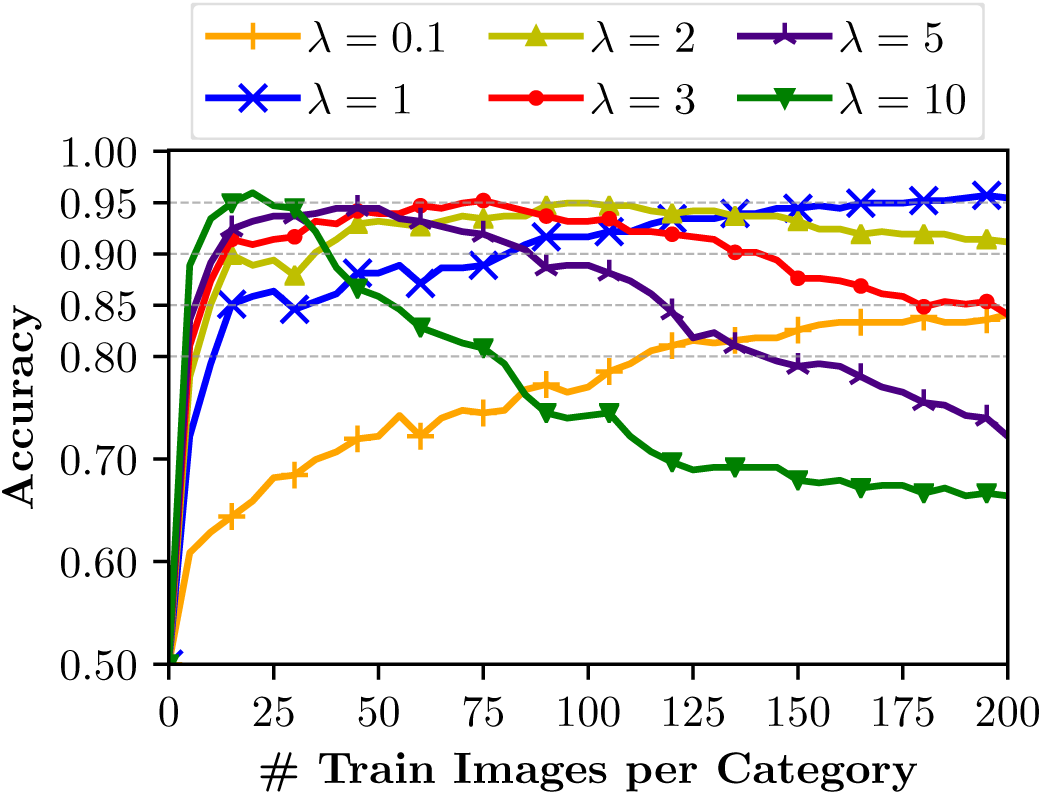
Impact of learning rate amplification on number of needed examples to learn. Accuracy on the test images as a function of the number of train images per category used in the co-occurrence Learning phase (increased number of shots in a few-shot learning task).

#### 4.2.3 Impact of the learning rate on few-shot learning

As anticipated earlier, we performed experiments to understand how the learning rates and number of examples shown in training interplay with each other. To give an intuition, we show Fig. 7 to illustrate the impact of the modulated (increased by some factor) learning rate on the accuracy as a function of the number of train images. Recall that the baseline is *α*^+^ = 0.007*, α^−^* = 0.003 and we multiply both values by some factor 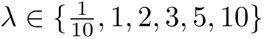. The figure shows the results for one of the train/test splits, picking the train images from the train set.

First, when *λ <* 1, the learning is obviously slower. Indeed, the accuracy varies from 50% with 0 training images (i.e. random), to 85% using the entire train dataset and grows in a linear way. It will probably reach higher values with more training time. When *λ* = 1, we reach the maximum ^19^ possible test score using all the train images.

Then, an interesting behavior emerges when *λ >* 1. The learning is at first faster, as it can reach high accuracy after having seen only a small sample of train images. However, it then starts decreasing as the number of train images increases. We recall that the used STDP rule keeps the weights within the range [0, 1]. Thus, starting with high values for *α*^+^ and *α^−^* allows to faster associate discriminant features with the images categories. However, as the number of train images increases, weights might become less helpful if they tend to reach the maximum value of 1 and to have thus smaller intermediate values. The scores would become closer and less distinguishable. Another factor is that, as we will see later, the SNN is better at recognizing and learning from certain particular images compared to others (see description of Fig. 8). Hence, being exposed to a good image with a higher learning rate leads to reaching a good accuracy. But being exposed afterwards to a “bad” image will lead to unlearning the good weights, thus decreasing the performance. When *λ* = 10 and with only 20 train images per category, it is possible to reach an accuracy of 95.5%. With *λ* = 2, 3, and 5, the number of necessary train images per category to reach the same accuracy are 45, 75, and 100, respectively.

**Fig 8.**
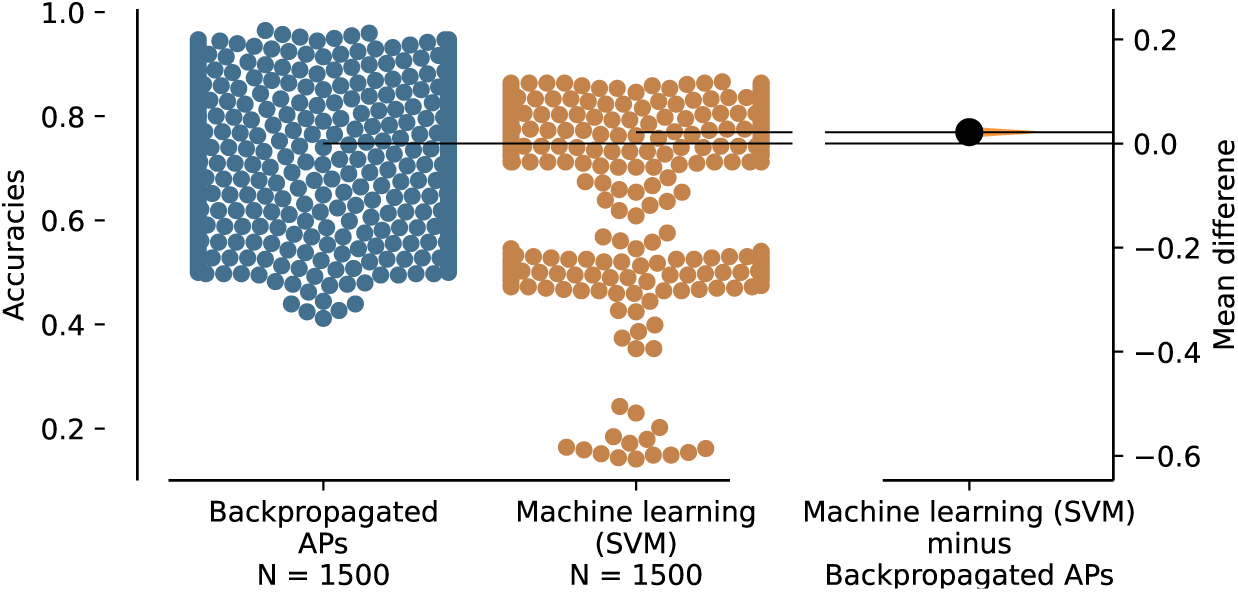
Comparison of ML and our hypothesis in one-shot learning tasks across 1500 experiments where each time, a single different pair of labeled (motor,face) images is shown during training. Evaluation is done on the usual test set of around 200 images. When using the backpropagation-based hypothesis, there exists specific (motor,face) examples that, shown to the SNN only once, yield maximum accuracy on the entire unseen test set.

These results hint to the following direction: the “best” approach in terms of few-shot learning would be to first start with a high learning rate or *λ* but then to stop changing the weights by either suddenly decreasing *λ*, or by simply freezing the learning and making the network always stick to the old “beliefs”. As earlier mentioned, this motivates the idea of using decaying learning rates, simulating an increased learning of novel stimuli and a progressive habituation afterwards, but this is out of the scope of the current paper. We hence leave it for future work. Here, we only use these results to understand the impact of *λ* and set it for the one-shot learning task. As anticipated earlier in the methods section, we set *λ* = 65 for our next last task.

#### 4.2.4 Task 2: Backpropagated-APs can outperform SVM in one-shot learning

In this more challenging task, the “teacher” shows the SNN only a pair of examples, one from each class. Fig. 8 plots [215] the Gardner-Altman estimation plot opposing the performances of our hypothesis and the SVM classifier on 1500 single pairs of examples each. First, despite the difficulty, on average, we find again that the backpropagation-based recollection hypothesis is still competitive with the SVM classifier (only 2%(95%*CI* : 1.2%*to*3.1%) less accuracy). Then, more interestingly, the performance highly depends on which particular couple of (motorbike,face) images was used for the one-shot learning task. We found that, *using the backpropagation-based recollection, there exists single couples of motorbike and face pictures that, whenever trained on, the model generalizes yielding an accuracy of 96.2% on all the remaining unseen* 198 test images. This is not only higher than what we achieve with SVM (maximum of 84.5% of accuracy), but even higher than what we achieved earlier when training with all images and not only one or few shots (there, we obtained 95.7% on test set for this particular split). Finally, the clear difference between the two swarm plots hints to a fundamental difference between the two computational strategies.

The observed curious disparity between pairs of images could be also due to a still unperfect feature extraction of current SNN models (e.g. not enough invariance and consistency across images). Further investigation ^20^ on the differences between these successful and less successful image couples could be useful, but is again outside the scope of this paper. Here, we demonstrate the computational efficiency of backpropagation-based recollection, and quantitatively compared it to a machine learning classifier on the same (good but not yet perfect) learned feature vectors.

To conclude, simulation results constitute another argument in favor of the backpropagation-based recollection hypothesis, supporting our call for arms for future investigations.

## 5 Efficiency in image reconstruction

Building on our hypothesis, we take a computational approach to assess its efficacy in reconstructing an image from sparse neural activations that distinctly characterize it. This mirrors the recollection of visual stimuli from the firing of source pointer neurons.

### 5.1 Materials and methods

Central to our hypothesis is the notion of backpropagated currents moving from post to presynaptic neurons, as a mechanism for reconstructing visual stimuli. To emulate this, we employed a popular neural network architecture from computer vision, AlexNet [5], to model the feedforward processing of visual information. *While not biologically plausible* in many ways as we discussed earlier, a pre-trained convolutional neural network bares some *computationally useful* similarities with humans’s visual processing [9, 216]. For what interests us computationally, they both (i) extract features of increasing complexity as we move deeper in the forward processing layers until they (ii) reach sparser representations that allow, for example, to linearly separate objects into distinct classes. We set out to reconstruct an entire image, using backpropagation, only from the sparse neuron that uniquely characterize the image once it is fed to AlexNet. If this procedure is successful, this would teach us that the backpropagation-based recollection principle is computationally sound.^21^

#### 5.1.1 AlexNet Architecture

Fig. 9 depicts the architecture of AlexNet, where several convolutional and fully connected layers are stacked serially, interspersed with “deterministic” max pooling operations that do not involve weight learning. Feedforward processing begins with the input image being properly prepared through resizing, center cropping, and normalization to conform to the input requirements of the pre-trained AlexNet architecture, a process encapsulated by the ForwardPrepare() function. The created tensor is then fed through the sequential layers of the network. Each convolutional layer applies learned filters, capturing hierarchical features ranging from edges in the early layers to complex patterns in deeper ones, activated by a rectified linear unit (ReLU) function (not shown in the figure) to introduce non-linearity. Max-pooling layers reduce spatial dimensions, providing robustness to translation and distortion. The sequential application of convolution, ReLU activation, and max-pooling operations encode the input image into progressively abstract representations as the data flows forward through the network. Finally, the feedforward pass concludes at the fully connected layers, culminating in a vector that embodies the high-level features extracted by AlexNet for the input image. This final vector of 1000 “neurons”, corresponds a probability distribution over the 1,000 class labels that AlexNet was originally trained to identify, with the classification outcome determined by simply identifying the index with the highest probability.

**Fig 9.**
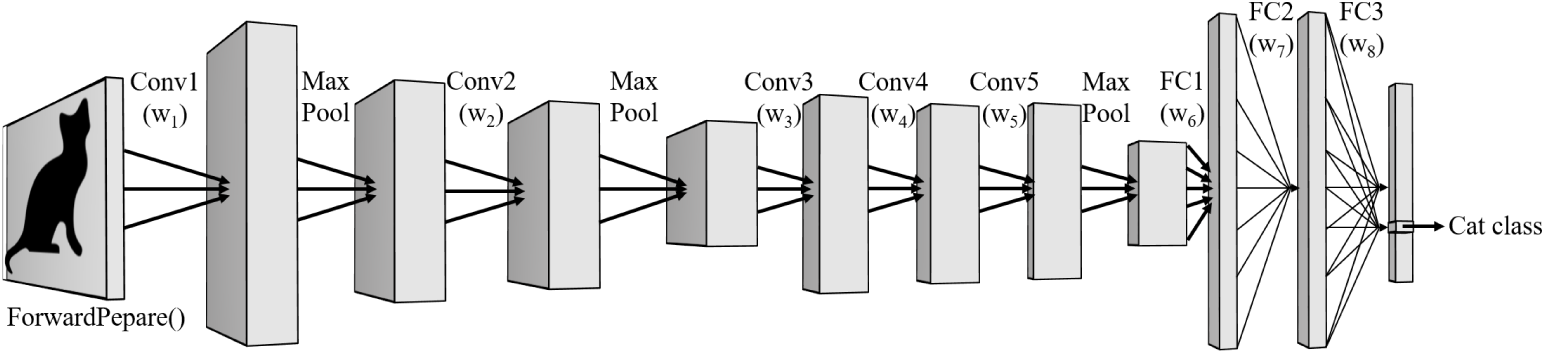
Schematized illustration of (forward) image processing with AlexNet.

It is exactly from the activation values of this last layer of 1000 neurons, i.e. the sparse neuron, referred to as *S*_0_ in Fig. 10 that we will attempt the first ambitious simulation of our backpropagation-based recollection. We also attempt reconstructions from the activations of intermediate layers *S*_1_ until *S*_10_.

**Fig 10.**
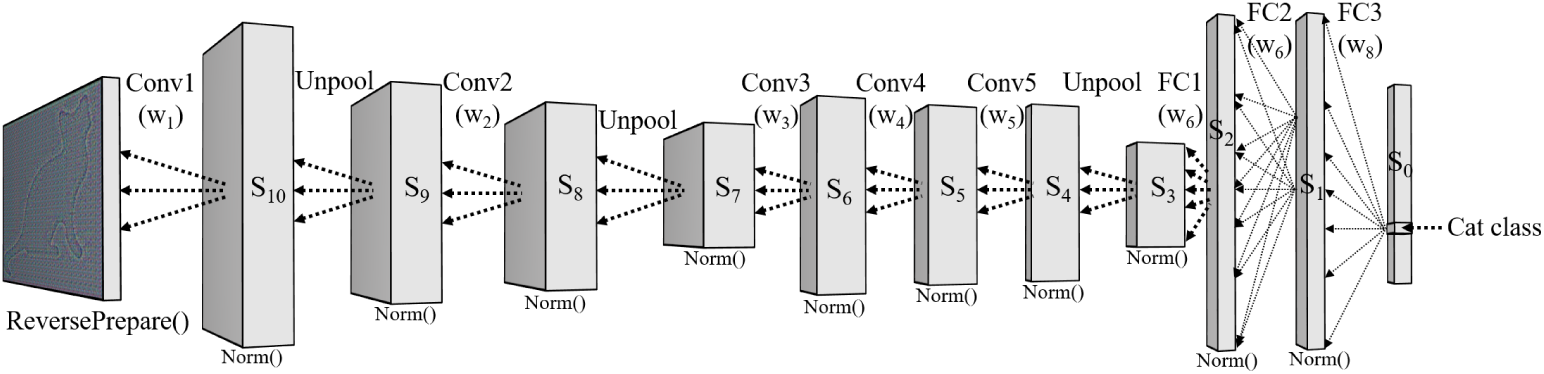
Backpropagation-based reconstruction on top of AlexNet.

#### 5.1.2 Backpropagation-based reconstruction on the AlexNet architecture

Upon completing a forward pass using the learned weights of AlexNet, we extract the 1000 last activations, which should best characterize the image. For our complete experiments, we also (i) use the argmax function to determine which one of these 1000 activations mirrors the classification output (ii) and store all the intermediate activation outputs, *S*_1_ til *S*_10_. We then initiate the backpropagation processing starting from the chosen source neurons, which we vary from the single sparse neuron that best characterizes the input, to the entire 1000 activations of *S*_0_, to each of *S*_1_ until *S*_10_. *Note that, of course, in each scenario, all the “pre-synaptic” activations are not used for the reconstruction process, only the weights are. This effectively mirrors a reconstruction process that happens during a later recollection event, based only on reactivating a set of sparse neurons that uniquely characterize the input*.

Fig. 10 illustrates the backpropagation-based recollection on top of AlexNet in reverse, throughout the layers. As can be seen there, at each step, we reconstruct the activations of the previous layer, by backpropagating the activations of the current layer, backwards proportionally to the corresponding weights. In doing so, three types of layers need to be traversed in reverse following the laws our hypothesis: dense layers, convolutional layers, and pooling layers.

##### Reconstruction on dense layers

Implementing backpropagation-based recollection on dense (i.e. fully connected) layers is easier, compared to convolutional layers, since each activation in the output is “connected” to all “neurons” in the previous layer, through a single weight matrix. Hence, given a current layer’s weights and its activations, we can derive the “reconstructed input” by performing a matrix multiplication of the activations with the transposed weight matrix. In essence, this is equivalent to distributing the activations backward through the corresponding weights, yielding a reconstruction of the activations of the previous layer that is faithful to our hypothesis: each neuron in the input layer receives the sum of “currents” from all neurons in the next layer proportional to the weights connecting both.

Fig. 11 exemplifies such a procedure through a toy example, with 9 input neurons at layer *ℒ*^0^ (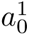 till 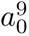), connected to 2 output neurons at layer *ℒ*^1^ (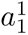 and 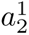), through 18 weights. During the forward pass, in deep learning terms (bias is omitted for simplicity), the output activations at *ℒ*^1^ are obtained via a matrix multiplication between the activations at *ℒ*^0^ and the weights matrix, which has a shape of 2 rows and 9 columns. During reconstruction, the activations of *ℒ*^1^ are backpropagated proportionally to the weights, and each neurons at *ℒ*^0^ receives contributions from each of the two neurons at *ℒ*^1^, which are simply summed.

**Fig 11.**
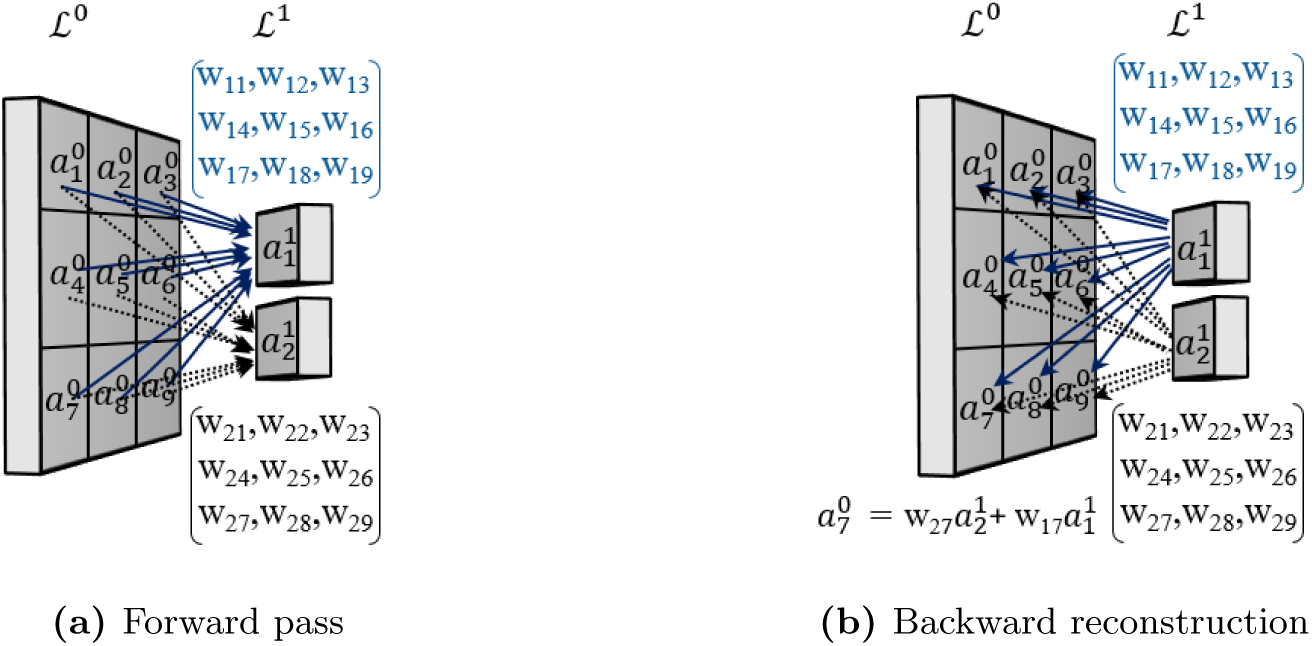
Toy example to visually illustrate backpropagation over the learned weights of AlexNet (The principle is valid for both Convolutional and fully connected settings).

In the end, computationally, such reconstruction procedure is equivalent to multiplying the activation matrix of the root layer (exemplified here with *ℒ*^1^) with the transposed weight matrix. This is further described in Algorithm. 1 in a self-explanatory manner.

#### Algorithm 1 Reconstruction of Input in Dense Layers with Optional Normalization

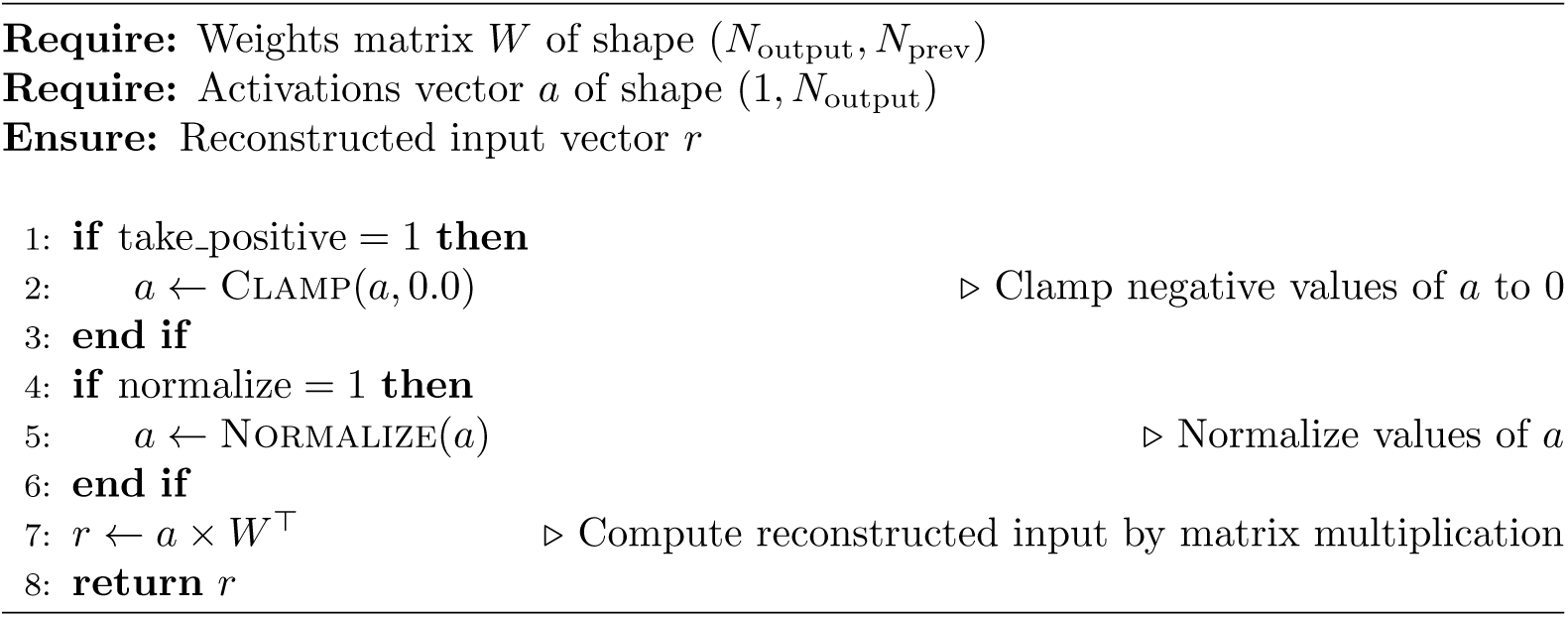

##### Normalization options

Once we apply the first reconstruction on the dense layer as just explained, we end up with a reconstructed activation matrix spanning a wide range of values.

As recently surveyed by Shen et al. [217], the brain employs a set of strategies to regulate neural activity paralleling several normalization techniques that are fundamental in contemporary deep neural networks. The authors identify four of such strategies in the brain and three corresponding equivalents in deep learning. First, (i) single neuron normalization adjusts individual neuron firing rates, ensuring a balance between excitatory and inhibitory signals, ensuring that the neuron maintains a target firing rate on average. This is similar to batch normalization in deep learning [218]. Second, (ii) synaptic scaling in the brain, adjusts the strength of connections between neurons, based on the neuron’s average firing rate, ensuring that all incoming excitatory synapses are appropriately scaled to maintain a target activity level. This in a way parallels weight normalization in deep learning [219]. Third, (iii) the equivalent of Layer-wise normalization in deep learning is known in the brain as divisive normalization [220, 221]. It involves adjusting the activity of a whole layer of neurons based on their summed activity. Finally, (iv) network-level normalization extends the principle to the overall brain activity. This is reflected by network homeostasis in the brain, where the overall firing rates and synaptic weights across a neural network are maintained stable despite local variability.

We accordingly experiment, during reconstruction, with few normalization options, focusing on the activations, mainly along the last two strategies.

- **Normalization** Our first option is simple Min-Max Normalization, which consists in scaling the reconstructed input to a [0, 1] range, somewhat representing firing rates, using the global minimum and maximum values. We perform two types of normalization, mirroring the last two strategies above: a local and a global one, and observe no major difference in reconstruction performance. The local normalization applies the operation locally to each channel or feature, using local minima and maxima, while the global one considers the entire input. Although, we obtain good reconstructions with this normalization, we experiment with other types of transformations for the sake of completeness.
- **Standardization** This option instead applies what is known as standardization which rescales the data to have a *mean* = 0 and *std* = 1. Similarly, we perform it either locally (per-channel or per-feature) or globally, and equally found no visual difference between global and local standardization, as performed equally well.
- **Rescaling** Finally, we also experimented with rescaling and clipping on top of standardization, using both local and global variations. Here, after standardization, either globally or locally, the data is rescaled to center around the value 0.5 and then clipped to ensure all values are within [0, 1].

Finally, we also tested the *option* to consider *only positive activations*, before normalizing, akin to the biological activation thresholds observed in real neurons, ensuring that only “excitatory” activations contribute to the reconstructed input.

##### Emulating backpropagating APs on Convolutional Layers

The methodology for simulating backpropagation-based recollection in convolutional layers involves a slightly more complex implementation due to weight sharing, where the same kernel—a small set of weights—is applied across the entire output in a sliding manner. But the principle that we exemplified in the toy example of Fig. 11 remains valid here too, and our objective the same: to reconstruct the input of the previous “pre-synaptic” layer (e.g. *ℒ*^0^) by tracing the reverse paths leading to the observed activations in the current layer (e.g. *ℒ*^1^). This is of course done using only the weights and activations of that current layer. The process involves calculating the contribution of each activation (e.g. at *ℒ*^1^) to its receptive field in the input (e.g. at *ℒ*^1^), followed by accumulating these contributions to form the reconstructed input. This procedure is better detailed in the pseudo-code outlined in Algorithm. 2. The full source code is also available.

##### The Winner-takes-all option

Another mechanism that is closely related to normalization in regulating neural activity is neural competition, where neurons compete for activation and only the “strongest” signals are propagated, while the others are inhibited. This is often embodied in the famous Winner-Takes-All mechanism [222], crucial in biological systems for tasks like feature detection and attention modulation. In the case of AlexNet, a close mechanism could implemented, but only in convolutional layers where different feature maps compete for the same spatial location. Hence, in addition to normalization, we experiment with adding some sort of winner takes all to Conv() layers.

The function, described in Algorithm. 3, operates on the activations *A* of a convolutional layer, structured in the typical format (*B, C*_out_*, H*_out_*, W*_out_), representing batch size, number of output channels or feature maps, and spatial dimensions, respectively. The core idea is to enforce a competitive environment where, for each spatial location, only the neuron with the maximum activation, across channels, is allowed to “fire”, while others are suppressed. This is achieved by initializing a zero-valued tensor *A^′^*of the same shape as *A*. For each batch and spatial location, the algorithm identifies the channel *c*_max_ with the highest activation. It then selectively retains this maximum activation in *A^′^*, setting all other channel activations at that location to zero.

#### Algorithm 2 Backpropagation Emulation for Convolutional Layers

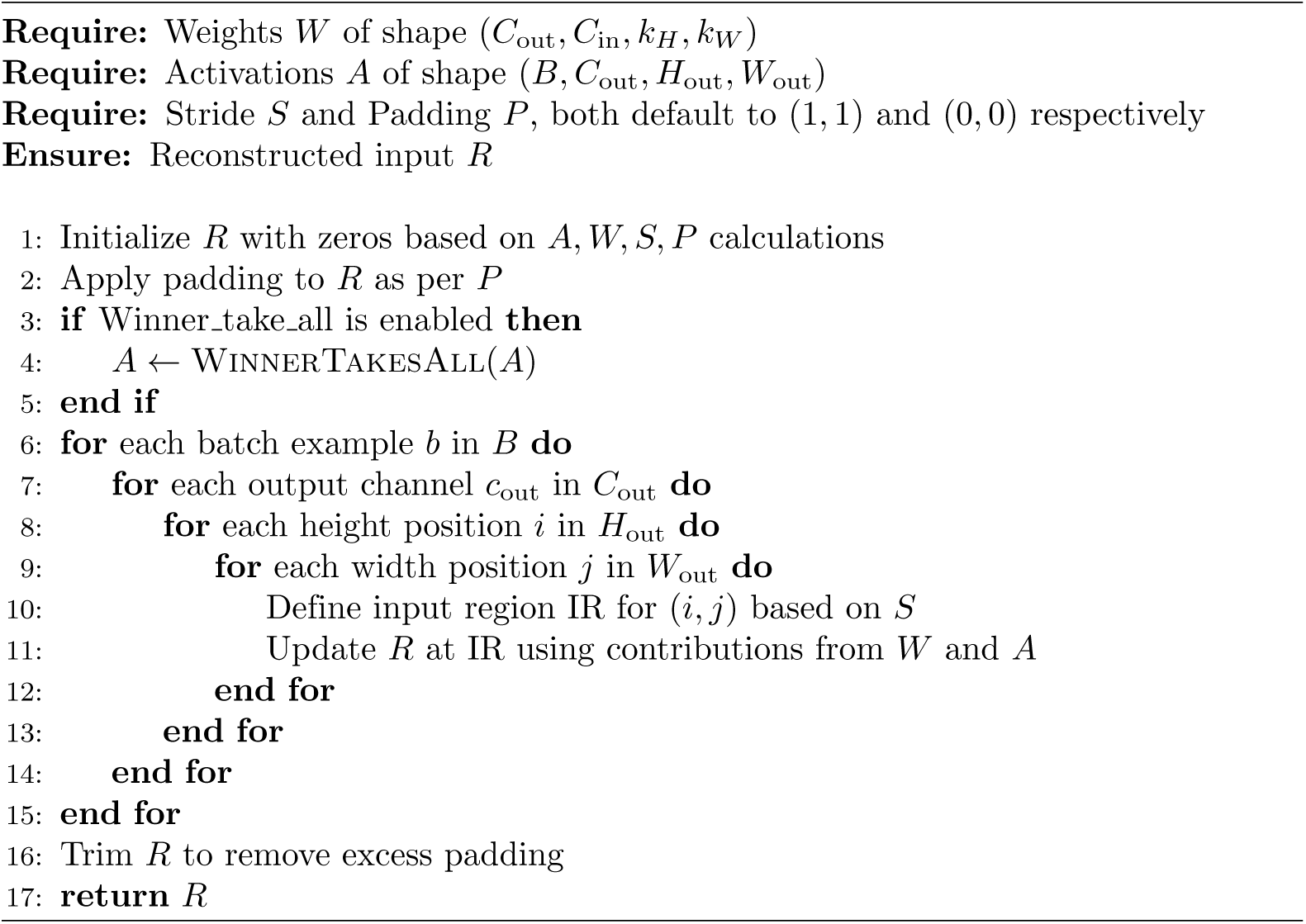

#### Algorithm 3 Winner Takes All Function

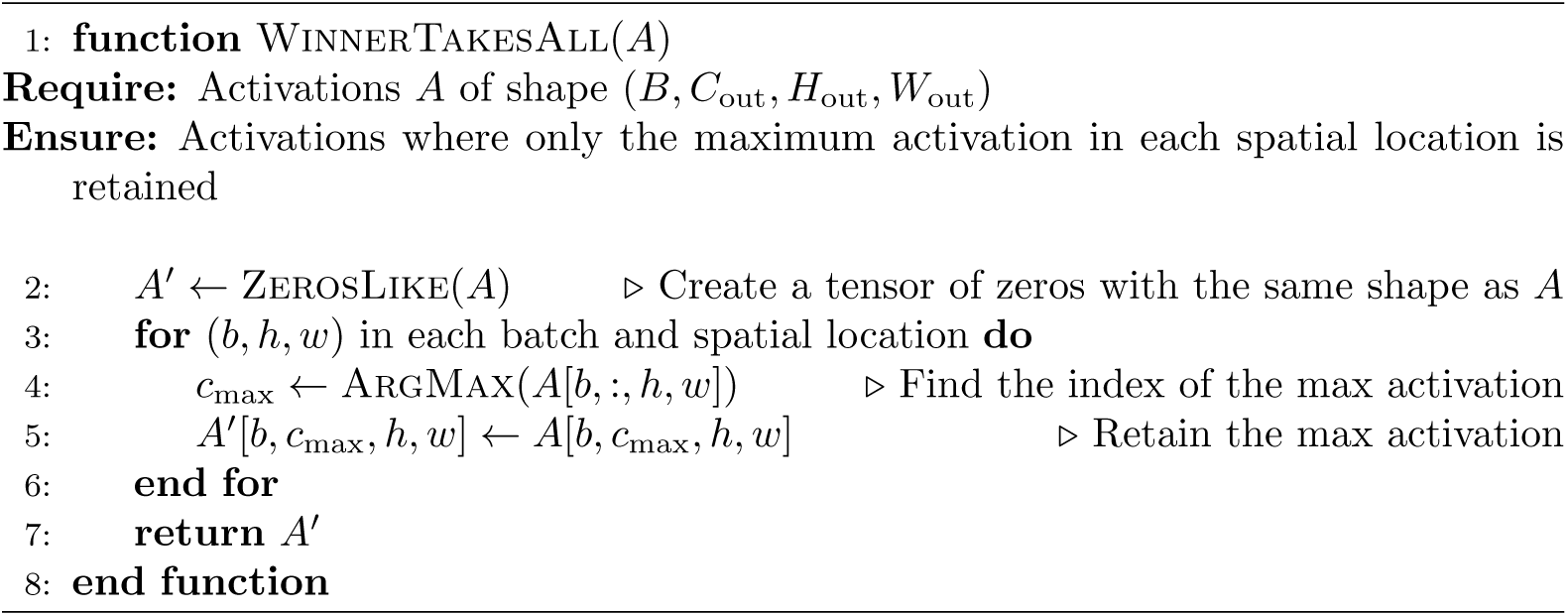

Note that a very similar mechanism has been called inter-map lateral inhibition in the work of Kheradpisheh et al. [3]. There, the authors used it during forward propagation of spikes in a convolutional neural network: when a neuron fires in a specific spatial location, it prevents other neurons in the same location, but belonging to other feature maps, from firing, effectively inhibiting them.

##### Reversing Max Pooling layers

Finally, *Max Pooling* layers do not use learned weights and are hence *by design impossible to be reconstructed* using backpropagated signals over the weights graph. Used in deep learning but to the best of our knowledge not biologically plausible, Max Pooling is a lossy operation. Indeed, as exemplified in the toy case of Fig. 12, once the maximum activation over a given region is computed, we obtain its value as the corresponding output, and discard the rest. In this operation, there is no associated weight to recover its location back. For this reason, we opt to reverse this operation in AlexNet deterministically, i.e. by resorting to PyTorch’s unpool() function, while retaining the indices responsible for the operations in the forward pass. Such unpooling is equivalent to a scenario where each neuron locally “remembers” which neurons in the previous layer led to the maximum activation, and backpropagate the signal only to those. Fig. 12 visually exemplifies the process in a self explanatory manner.

**Fig 12.**
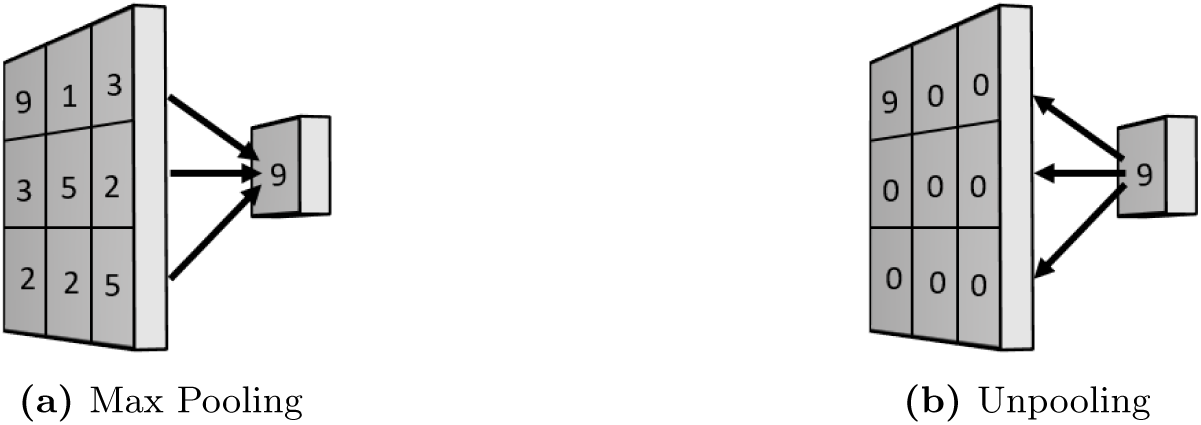
Toy example to visually illustrate the Max Pooling and Unpooling operations we employ in AlexNet.

Note that this inevitable choice leads us to keeping some state from the forward pass, i.e. the indices of the neurons that led to the max operations in the pooling layers. Nonetheless, although pooling layers constitute only 3 of the 11 layers that we traverse, our first results showed that this choice had drastic and surprisingly positive effects on the performance of the reconstruction. This finding lead us to looking for alternative architectures that do not employ pooling layers. To our surprise, the literature fell short in providing such pre-trained language models, leading us to train a custom pooling-free architecture in Sec. 5.1.3.

`BackwardPrepare()` **options** But before that, once the reconstruction done, we need to visualize the resulting image to qualitatively evaluate the reconstruction. Here, we experiment with three simple options. The first is the exact reverse of `ForwardPrepare()`, since the latter is fully reversible. The second is standardization and rescaling, and the third is to do nothing but the default min-max normalization.

#### 5.1.3 Backpropagation on a Pooling-free AlexNet-like architecture

Next, given our curious findings on the role of Max Pooling which we will detail later in the results, and in the absence, to the best of our knowledge, of a pooling-free pre-trained neural network for image classification, we opted, for the sake of completeness, for retraining a custom model. For this, we start from AlexNet and perform slight modifications. In particular, we replace its pooling layers with equivalent strided convolutions of similar shape. The latter result in the same amount of downsampling, but using learnable weights, instead of the deterministic rule of max pooling.

When building our custom model, we inherit the pretrained weights of AlexNet, and start from random weights for the newly added strided convolutions. We then experiment with two types of training which yielded similar results. In the first, we train only the new strided convolutional layers, and freeze the rest. In the second, we gradually retrain all the layers, first training only the pooling replacements, and slowly unfreezing the rest of the network. Both approaches yielded similar results.

As for training itself, we do it on a classification problem using the ImagenetV2 dataset [223], which we split into train and validation. We employ the categorical cross entropy as loss. Noticing that we failed to learn an accurate classifier without pooling layers^22^ (perhaps explaining why computer vision architectures still need them), we compared two training strategies. In the first, we pick the model that performs best on both training and validation set, as per classic machine learning good practice. In the second, we let the model overfit on the training dataset, even if it performs poorly on the validation. This is a reasonable choice as our goal is *not* to reach the best classification performance but to understand the behavior of backpropagation-based recollection, as a function of the learned weights.

This experiment complements the main AlexNet one as follows. Unlike AlexNet, which has good quality weights as witnessed by its classification performance, our custom model is not a perfect classifier, so we expect its reconstruction to be of poorer quality. However, it has the merit of having only learnable weights and no deterministic pooling layers. This allows us to demonstrate two key aspects of our hypothesis: (i) the specificity of the reconstruction based only on the activations resulting from the forward pass and (ii) the effect of sparsity on the reconstruction quality.

## Code Accessibility

The datasets and code for this study will be found in the same git repository as the previous experiments: https://github.com/bendiogene/recollection_hypothesis.

### 5.2 Results

#### 5.2.1 Original AlexNet

We first show in Fig. 13 examples of original images, and their corresponding reconstruction using our method, starting from *S*_0_ on the original AlexNet with MaxPooling. There, we show the case where all the reconstructed activations, including the final BackwardPrepare(), undergo simple min-max normalizations. As can be seen, the images are nicely reconstructed, perhaps anecdotally, in a non-vivid manner that is reminiscent to how we actually recollect previously seen images.

**Fig 13.**
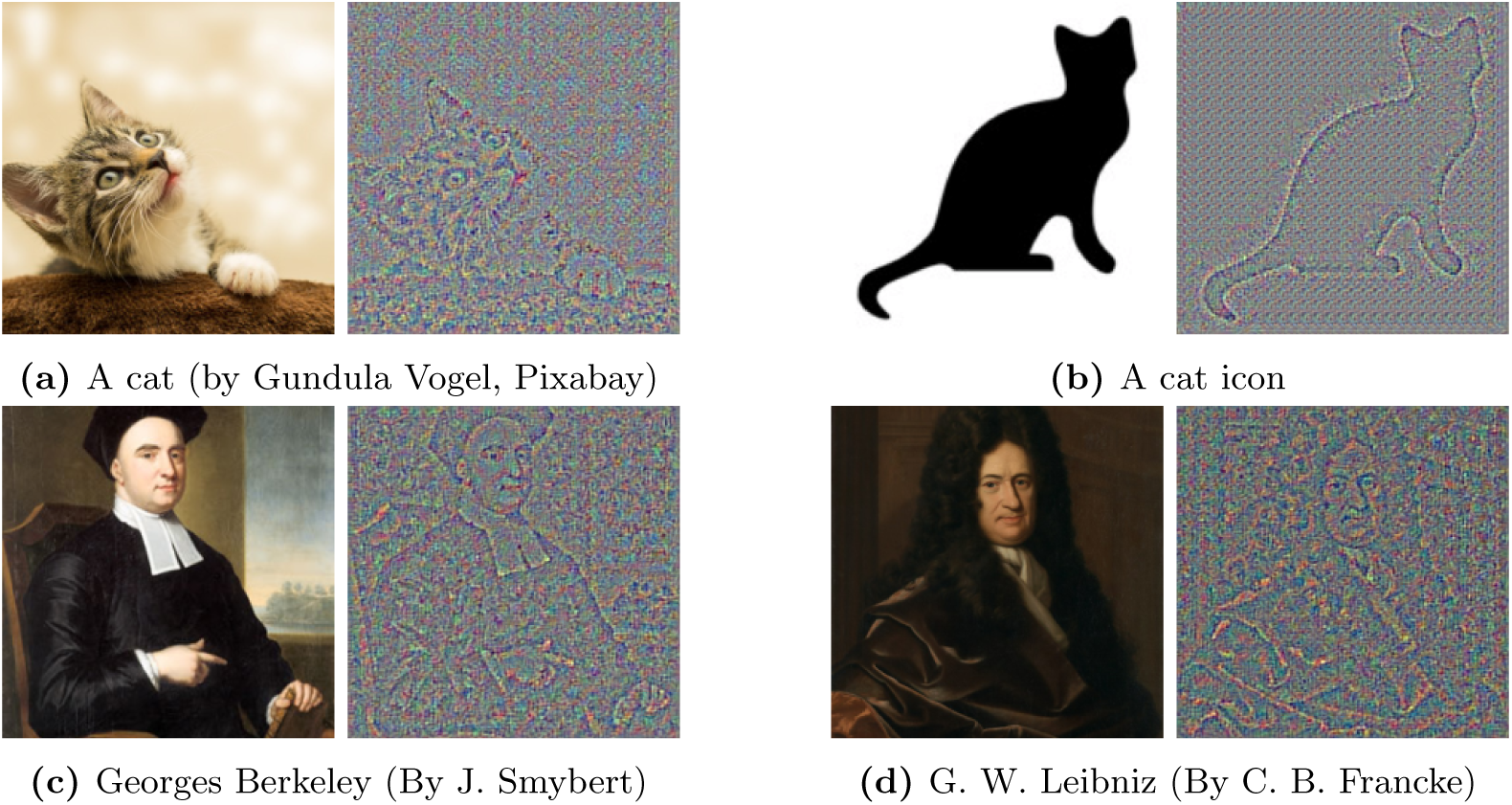
Comparison of original (left) and reconstructed (right) images

Next, we present the results of a series of experiments to understand the reconstruction process and assess the impact of various “design choices” on the quality of reconstructed images.

##### Necessity of both Unpooling and Backpropagation over weight graph

The first result is a sanity-check to verify the necessity of having backpropagation over both (i) weights and (ii) unpooling layers in order for the reconstruction to be successful.

Not illustrated with a figure, we found that randomizing either of the pooling indices or the weights leads to the failure of the reconstruction, demonstrating that *both unpooling following the max indices and backpropgation over weights are necessary for the reconstruction to be successful*.

##### Impact of Source Pointer Neurons and the surprising effectiveness of unpooling

Next, as the reconstruction from all activations in *S*_0_ worked already so well, we attempted to reconstruct the image from the single (top 1) neuron activation in the last layer (which also represents the class of the image). To our big surprise, this gave almost the same results as using all of the 1000 activations of the last layer. Not shown in Fig. 14, but will be released with our code, we verified that the reconstructed matrices are not identical, although the visual impressions are very similar.

**Fig 14.**
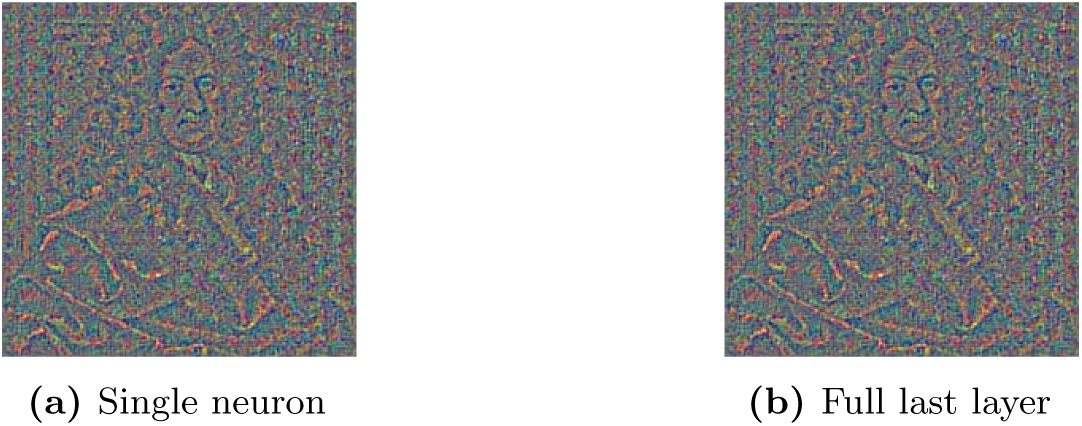
Source Pointer Neurons: Single Neuron vs. Last layer

However, what seems rightly unreasonable at first sight with this result, having in mind our hypothesis, is that this single source neuron does not only “represent” the specific image we showed to the network, but is supposed to signal all the images that belong to the same ImageNet class. But as we mentioned earlier, we do keep some state from the forward pass besides the weights: *the max pooling indices necessary to backpropagatedly recover the max pooling operation*. It turns out that it is this input that encodes the last memory, allowing to faithfully reconstruct it in particular, and not another one.

With further experiments, it turns out that these unpooling indices are so powerful that they help recovering the image even starting from noisy input. Not shown in a figure, the first experiment that has put such behavior in evidence was attempting the reconstruction using random activations at *S*_0_, and noticing to our surprise how the reconstruction was not affected. In fact, as partly illustrated in Fig. 15, this behavior continues to manifest itself when starting with random activations at *S*_1_, and continues to hold true up to starting from random activations at *S*_8_.

**Fig 15.**
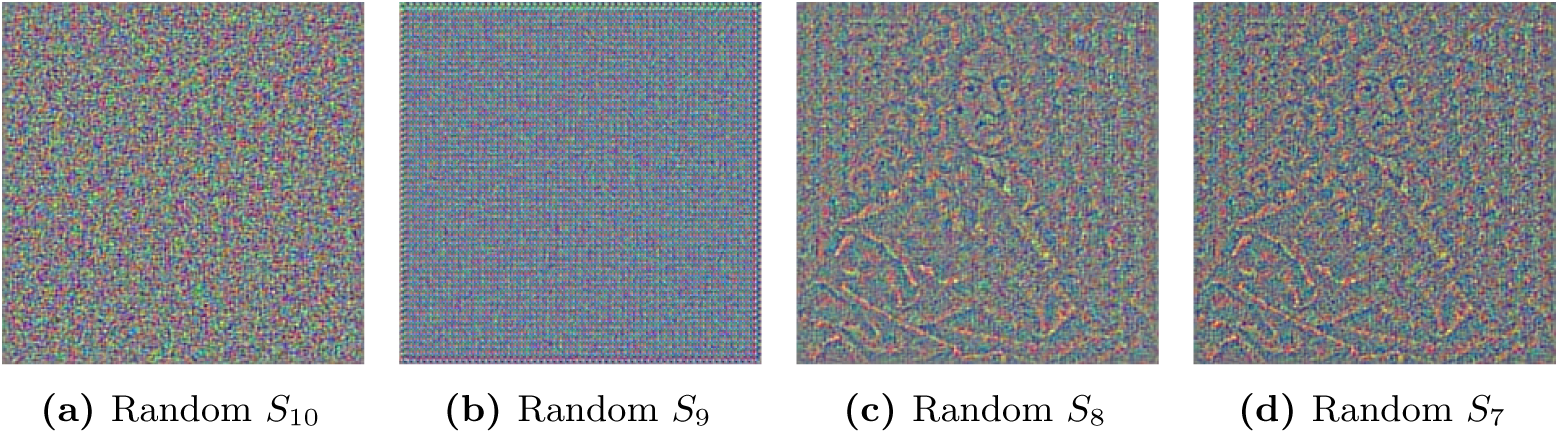
Illustration of the reconstruction power in the pooling-based AlexNet even starting from noisy activations.

The reconstruction from noisy (random) inputs becomes however impossible beyond: starting with random activations at either *S*_9_ (just before the last unpooling in the chain) or *S*_10_ (just before *Conv*_1_) fails to reconstruct the image, showing again, that both convolutional layer weights and unpooling indices are necessary for backward reconstruction: the unpooling indices are necessary to recover the particular image we showed in the forward pass, and the learned weights are necessary to make a successful reconstruction.

Hence, a crucial consideration here is that unpooling indices extract enough information from the forward pass to allow the recovery of the image, even starting from noisy inputs. This departs from the hypothesis that backpropagated reconstruction happens starting from *specific* activations and not from random inputs. To evaluate this particular aspect of activation specificity and sparsity, we study in Sec. 5.2.2 backpropagated recollection in our custom pooling-free neural network.

Nonetheless, AlexNet experiments represent a strong argument in favour of the efficiency of backpropagated recollection as a computational principle: both unpooling and backpropagation over weights are *necessary* for this reconstruction, and both use *local rules* where each “neuron” looks only to its presynaptic neighbors in order to backpropagate the reconstruction signal. The abilities of Max Pooling are however intriguing and merit further exploration in future work, as this is beyond the scope of our current hypothesis.

##### Effect of normalization

Next, a crucial function on which we rely in our simulation of the backpropagation-based recollection is normalization. For the sake of completeness, we qualitatively compare the impact of the various normalization approaches on the quality of the reconstructions in Fig. 16.

**Fig 16.**
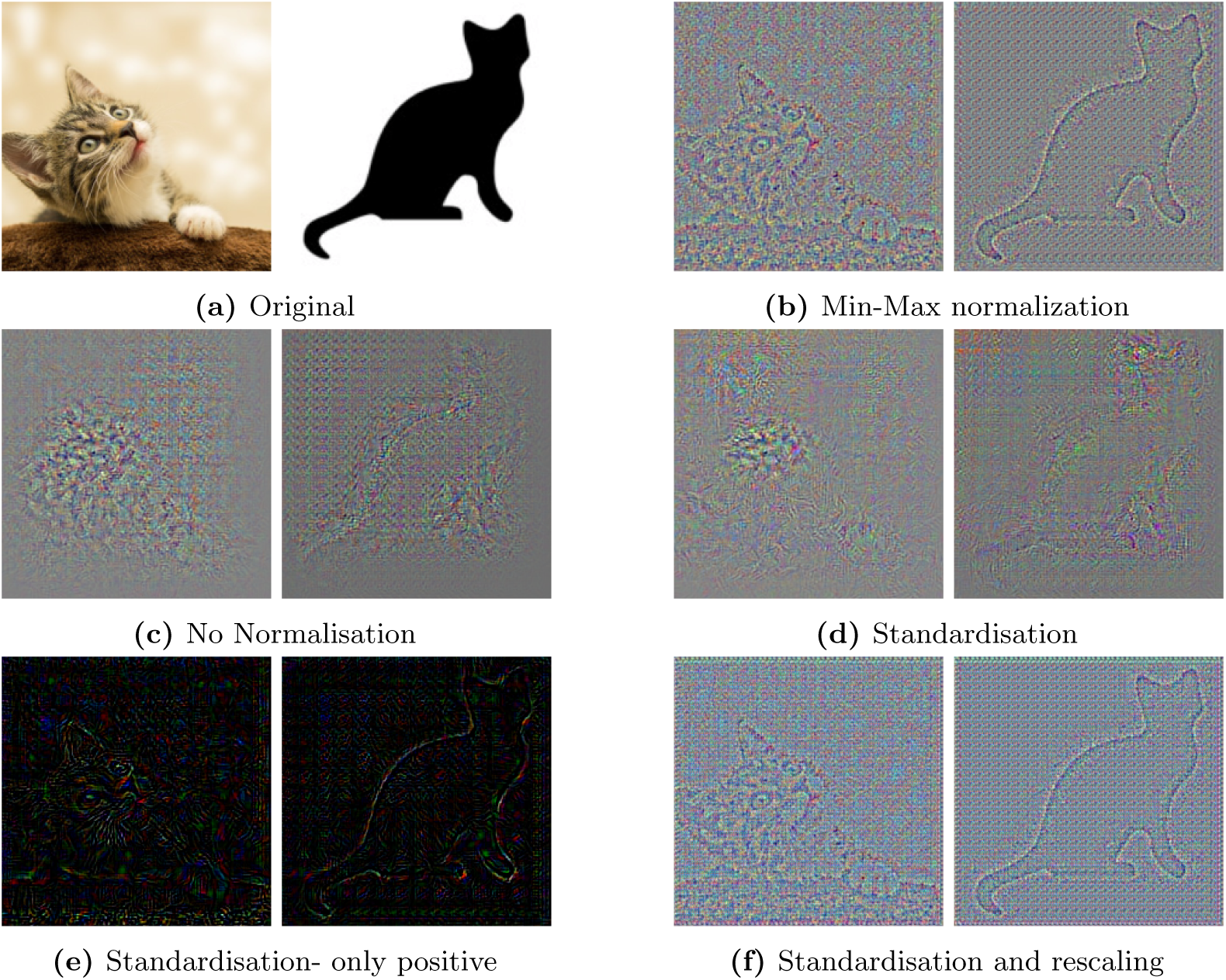
Impact of different normalization methods

As can be observed on the figure, Min Max normalization yields the clearer results. Standardisation alone without rescaling is not sufficient to have a good reconstruction, unless only positive activations are considered. Another observation is that applying rescaling after standardisation brings the same benefits as min max normalization, yielding similar good reconstruction quality.

Finally, and perhaps most importantly, not normalizing the intermediate activations leads to quite blurry reconstructions where the original image is only slightly perceptible. This is inline with the crucial [217] role of normalization in both the human brain and deep learning algorithms as we discussed thoroughly in Sec. 5.1.2.

Finally, we exemplify in Fig 17 the impact of the BackwardPrepare() functions. As can be observed, the three functions yield clear reconstructions. The reconstruction using the exact reverse of ForwardPrepare() is however visually slightly different.

**Fig 17.**
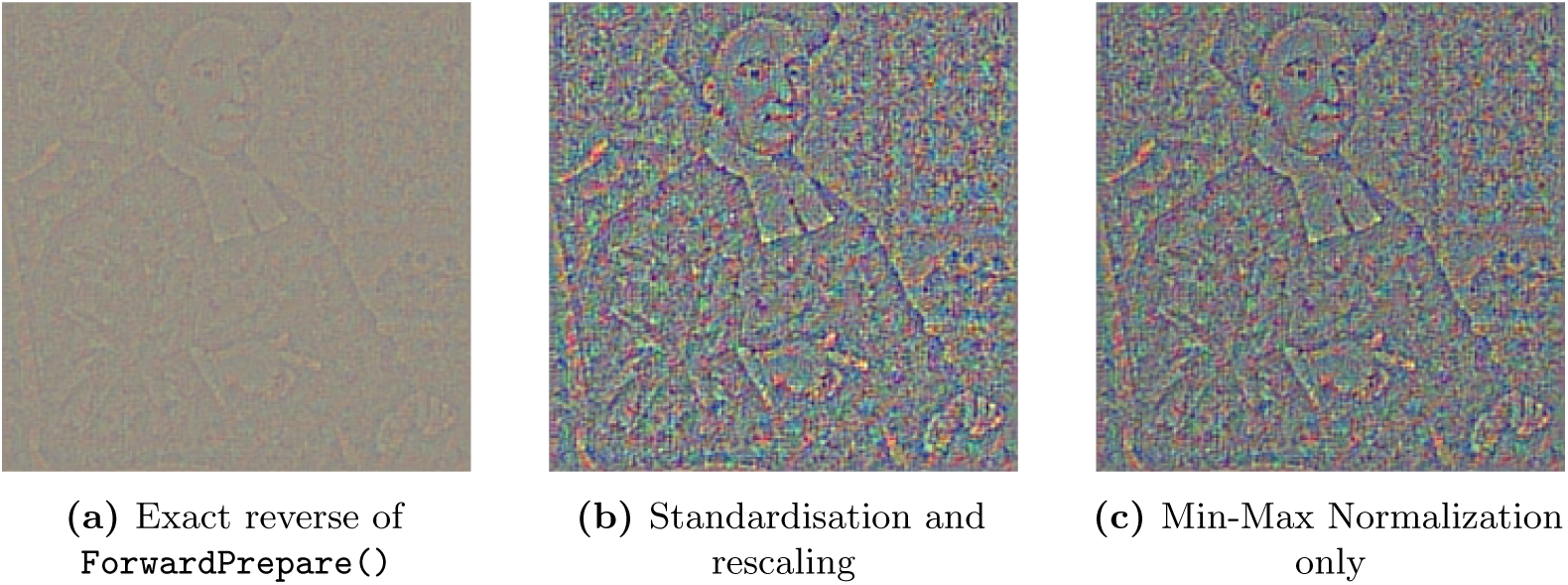
Comparison of BackwardPrepare() functions

We conclude that for what concerns AlexNet, the normalization method is crucial to the performance of the memory reconstruction.

##### Winner takes all to replace normalization

Finally, another way to biologically implement some sort of local normalization is lateral inhibition and the winner-takes-all mechanism. To isolate its impact, we start from no normalization at all, and activate for convolutional layers a winner-takes-all mechanism as explained in Algorithm. 3. We show the results in Fig. 18. While non normalized intermediate activations lead to bad reconstruction results as we saw earlier, simply adding the winner takes all mechanism leads to observably better reconstruction.

**Fig 18.**
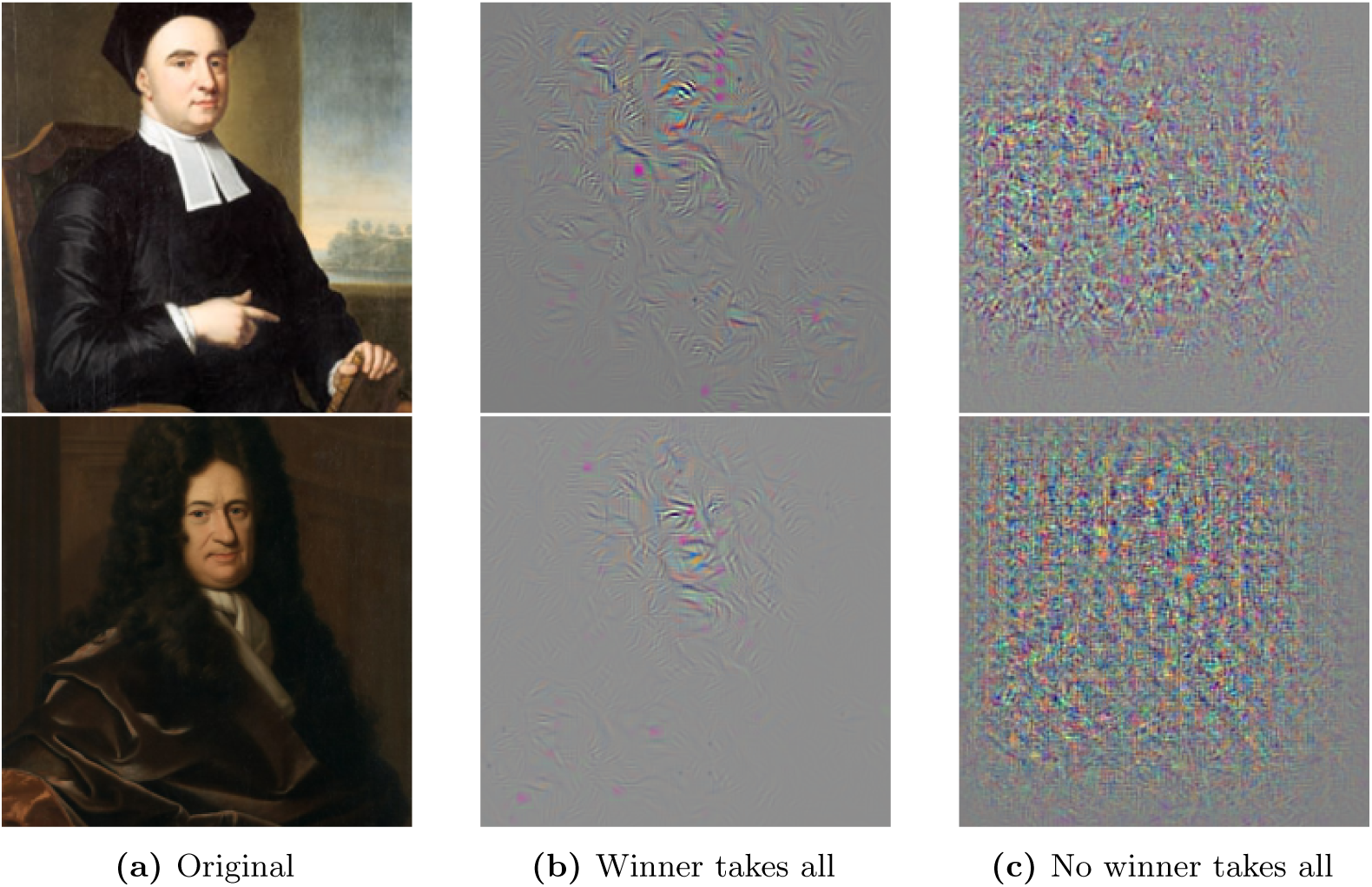
Impact of *winner takes all* strategy in case of *no normalization*

##### Takeaway

We conclude that backpropagation-based recollection on the AlexNet architecture yields good quality reconstructions, and that normalization is crucial for reconstruction. However, our use of some image-specific state from the forward pass (i.e. the pooling indices) makes that we cannot test a major aspect of our hypothesis: the ability to recollect an image *only from its feed-forward activations*. Indeed, the pooling layers of AlexNet, together with the learned weights, allow the recovery of the image even starting from random source activations. We next evaluate reconstruction on the pooling-free modified AlexNet using only learned weights and no pooling indices.

#### 5.2.2 Pooling-free modified AlexNet

Although we failed to train the Pooling-free AlexNet to reach a good performance compared to its pooling-empowered counter-part, we managed to train it in a best-effort manner, opposing two training strategies: the first is to overfit on the training set, and the second is to pick the best possible model that still performs well on the validation set, as per machine learning best-practice.

The results of these two experiments, when retraining only pooling replacements, are depicted in Fig. 19 and Fig. 20 respectively. Needless to remind but random weights lead to the *incapacity* of the network to reconstruct images, as also shown earlier. Both figures show the reconstructed image as a function of the root layer.

**Fig 19.**
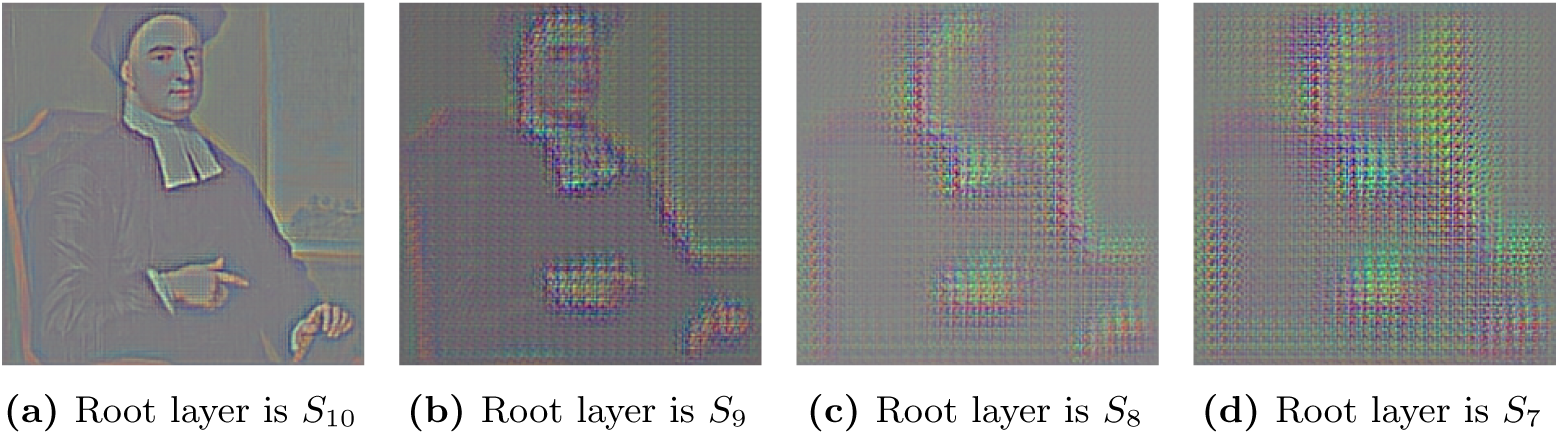
Reconstructed images as a function of source root layers (Pooling-free AlexNet - Retrain Pool only - Overfit on training set)

**Fig 20.**
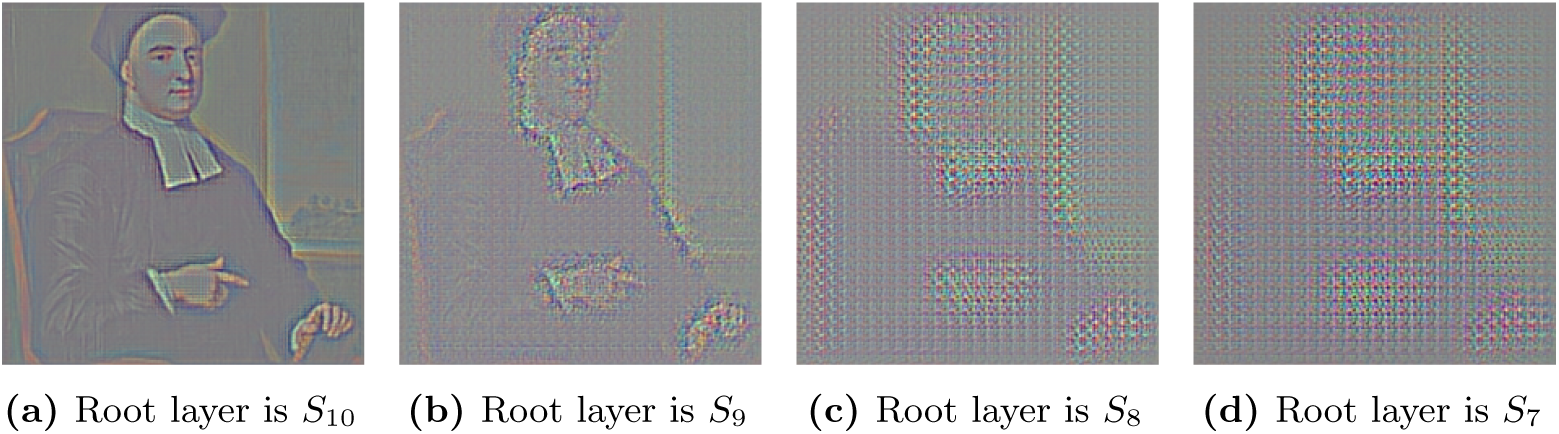
Reconstructed images as a function of source root layers (Pooling-free AlexNet - Retrain Pool only - Best classifier with no overfitting)

First, let us observe that reconstructing starting from *S*_10_ yields a nicely reconstructed image. This is expected as this reconstruction uses the original good quality AlexNet weights of *Conv*1.^23^

Reconstructing the image starting from *S*_9_ is thus more interesting to analyze since the backpropagated signals would have to go also through the newstrided convolution replacements of the old AlexNet pooling layers. There, two interesting observations can be extracted. First, the reconstruction is blurrier, compared to starting from *S*_10_, but is still feasible as the silhouette is clearly visible (again, compared to random weights which, we remind, give random reconstructed pixels). Second, the reconstruction using the “best classifier” yields higher details compared to over-fitting. This clearly suggests that the better the Neural Network is at its tasks, and hence the better the learned representations, the better is the reconstruction. This observation is particularly powerful, since the neural network’s training objective was only classification, and of course, not reconstruction. We verified that this behavior is consistent across 10 different training runs (5 with overfitting and 5 with best-classifier).

Finally, the deeper we go in the layers, the more difficult the reconstruction becomes. This is not surprising, as we did not succeed in learning a good classifier, and as the errors introduced by each layer probably add up to worsen the quality of the reconstruction.

##### Impact of using sparse neurons

In our final experiment, we explore the influence of neuron sparsity on the quality of image reconstruction. Our hypothesis suggests that memory recollection is driven by a small number of highly active neurons, which initiate a backward flow of signals. Typically, less active neurons would not fire and hence would not contribute significantly to this process. This principle of neuronal sparsity aligns with the increasingly abstract representations of stimuli seen in both biological and artificial neural networks, though it’s not a unique feature of our backpropagation-based recollection hypothesis.

To understand this better, in the context of our theory, we specifically examine how the number of active neurons, or their “sparsity”, in a layer affects the reconstruction of an image. For this purpose, we use the layer *S*_9_ from our modified AlexNet model and selectively activate only the top N neurons based on their activity level. All other neurons are set to zero activity before we begin the backward reconstruction process.

Figure 21 showcases the results of this approach, presenting the reconstructed images that emerge from varying degrees of neuronal sparsity in the source layer. The figures shows two images, one simple, the icon of a cat, and the other with more details, the face of Berkeley.

**Fig 21.**
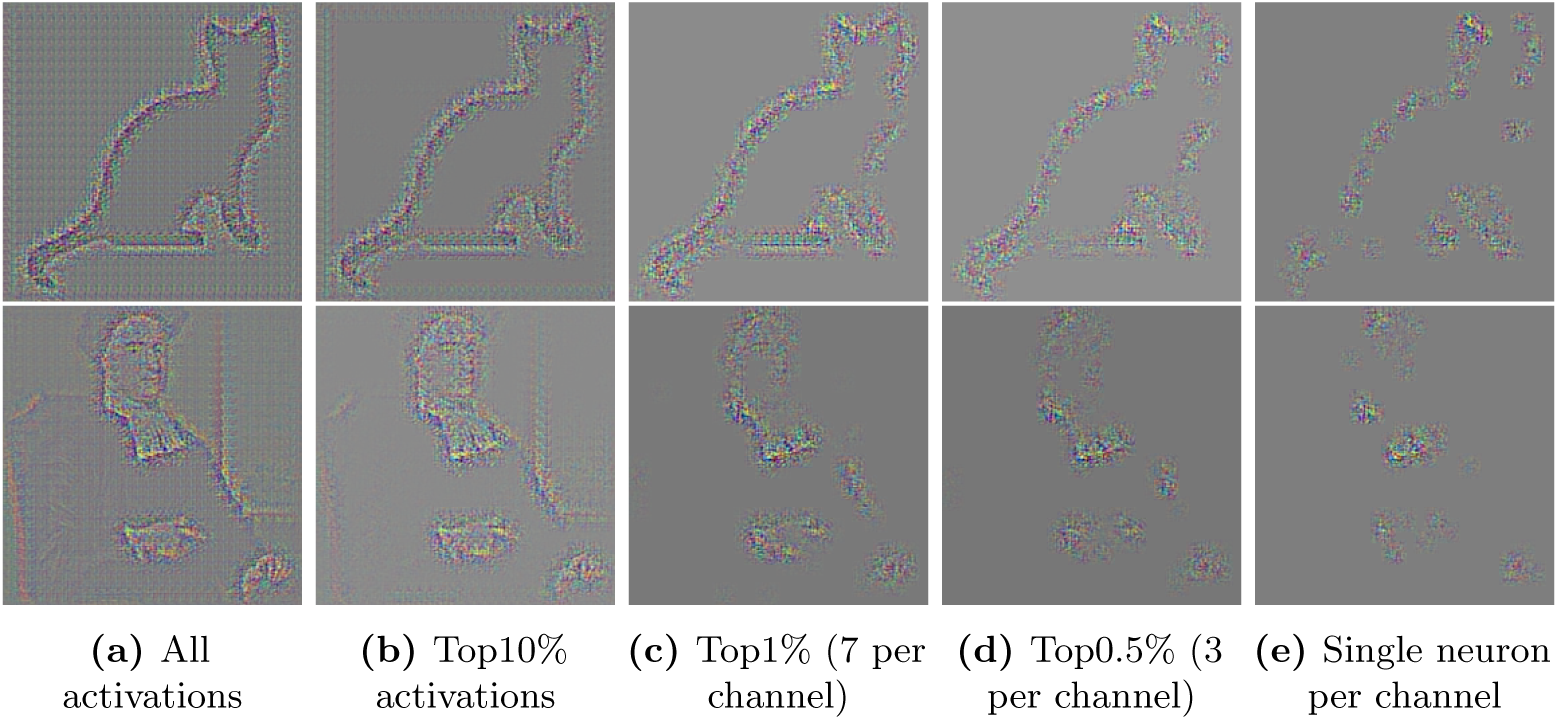
Sparsity. Reconstructed images from root layer *S*_9_ using decreasing number of source neurons

It turns out interestingly, that for the custom AlexNet (which, we remind, is not even a good classifier), starting from top activations leads to a recognizable reconstruction. In the case of the simple cat icon, taking a single neuron from each channel 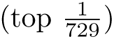 yields a distinguishable silhouette. This is a bit more difficult for the face of Berkeley, where it’s only slightly distinguishable with 7 neurons per channel.

The brain is of course a better “classifier” than AlexNet, not to talk about our custom AlexNet. We features of much better quality, the sparsity effect should be more manifest.

##### Takeaway

Unlike AlexNet, our custom model is not a perfect classifier, but it has the merit of having only learnable weights and no deterministic pooling layers. This allowed us to demonstrate two key aspects of our hypothesis. The first is that *reconstruction can be performed uniquely based on the activation pattern* produced in the feedforward pass. The second is *sparsity*, as we showed that neurons with the highest activations (i.e. those that would fire in response to a stimuli) are able to recover the image trace. Again, AlextNet is of course not a biologically plausible neural network, but similar pre-trained models are today the best available proxies to evaluate the efficiency of backpropagation-based recollection as a computational principle.

## 6 Discussion

For more than a century [224], information processing in the brain has been mainly believed to follow the forward pre to post-synaptic neurons direction. In this work, we emitted and evaluated the hypothesis that the backpropagation of APs in the reverse direction mediates many “offline” generative tasks where the simultaneous activation of specific targeted populations of neurons is needed. Our literature review shows abundant and intriguing evidence in favour of our claims (see summary in Sec. 3.6). As an added bonus, we computationally showed in Sec. 4 that backpropagated APs can be better than a machine learning algorithm in retrieving the name of an object. We further showed in Sec. 5 that image reconstruction is possible with backpropagation-based recollection. If confirmed true, our hypothesis will have tremendous implications.

### 6.1 Implications

The first implication is the promise to solve the neural encoding problem, reconciling the views of localist and distributed representation theories. In our framework, the answer to the representation problem becomes simple. First, (i) representations of concepts are hierarchically encoded (from complex to simple) by populations of neurons. At the same time, (ii) there exists highly selective source pointer neurons that respond to unique concepts; *their role is not to encode the concept but to allow the retrieval of its distributed representation or features*. These neurons act as hubs between various related encoded concepts, allowing the brain to “navigate” through them. For example, few neurons selectively respond to the image of a cat but they serve as pointers or hubs to easily connect such image to its related memories. Some of such neurons act as pointers to retrieve the visual features of a cat through backpropagated APs that selectively reactivate the neurons that represent the right lines, shapes^24^ and colors that define a cat. Hence, to be able to picture the image of a cat, the brain does not only need to activate the sparse neurons that respond to “cat”, but also, as confirmed by optogenetic studies above, entire populations of neurons. And if neurons that represent, say, vertical lines, are not activated during this process, the recalled mental image will lack them.

Beyond vision, the cat’s toy example could apply to any mental state and couple of *(lived stimuli, later memory of that stimuli)*, let them be smells, affects or even impressions of movements. In accordance with the principle of grounded cognition [225], for which we believe our framework applies, discrete concepts are grounded in the sensorimotor experiences that were encoded with them, such that the activation of a signifier of a concept leads to the activation of the experiences that are grounded with it. Hence, the cue that is the words “moving” or “tickling” is correlated with areas that encode actual moving or tickling.

Next, another hard problem on which our hypothesis could shed light is the binding one, i.e. how the brain binds higher-level concepts to more elementary ones or how it associates the right features (e.g. colors) to the right object if an image is composed of many. The fact that the brain needs some iterations and time to correctly perform the binding [226] suggests that this operation is not forward-based but happens iteratively later in a second stage. In our framework, this should happen through repetitive runs, involving top-down backpropagated APs from the right source pointer neurons of a single object, all the way backwards, to activate the neurons that encode its attributes.

In fact, two famous competing theories on this problem are (still) the feature-integration theory [227] and temporal synchronization theory [228, 229]. Our hypothesis promises again to reconcile them. Both theories admit the involvement of different runs of bottom up and top down hierarchies (e.g. attention in feature integration theory). However, the exact physical mechanisms are still unknown today. Our hypothesis obviously constitutes a perfect candidate. Moreover, under this new realm, one does not have anymore to chose exclusively between binding-by-synchrony and feature-integration theory: *attention* with backpropagated APs could *synchronously* activate selectively all the features related to a given discrete object.

An implicit consequence of this is that attention itself is likely implemented through top-down backpropagated APs that selectively reactivate the right distributed networks to attend to. Interestingly, evidence suggests that transient cholinergic modulation, which as we showed earlier triggers backpropagated APs, causally mediates cue-detection and attentional control, i.e. helping to focus attention on specific cues in a top-down manner [230, 199]. This can be seen as an immediate form of recall, where the cue related feature are recollected while the cue stimuli is still present.

In general, backpropagation could constitute an ubiquitous and easy-to-implement simple mechanism that underlies a diverse set of cognitive tasks such as offline thinking or mind wandering, imagination, episodic memory retrieval and future episodic thinking. In our framework, imagination would be “simply” the resulting activation patterns of a mixture of usually unrelated concepts: for example, an imagined “laughing cat” results from simultaneously back-activating the “laughing” and “cat” concept neurons. The same applies for mind wandering, where backpropagated APs should induce activation patterns that follow the most likely neural pathways, thus generating what seems to be coherent thoughts. Interestingly, all the above-mentioned tasks elicit subjective experiences that are transient and way less vivid than perception, a fact that could be explained by the weak and fading-away nature of backpropagated signals.

Moreover, our hypothesis could open new ways to better understand some recollection-related pathologies. In this case, understanding what factors impact the inhibition or excitation of backpropagated APs could help understanding probably-related disorders such as obsessive thinking or intrusive thoughts. Similar (rarer) dysfunctions happen in mental visual imagery as well, such as the absence (aphantasia) or excess (hyperphantasia) of visual imagery experiences [70]. In all these cases, the neuromodulation mechanisms that control backpropagation should be first suspected.

Last and not least, our hypothesis implies that the retrieval process is stochastic in nature: the retrieved memory trace looks like the original first perception but, depending on past experiences (and hence the weights of the neural connections due to past experiences), the reactivation might not be exactly the same. This can be seen most in the case of language where the same word (e.g. “a screen”) was seen multiple times during encoding, in the presence of multiple similar stimuli (e.g. many different types of screens), and where the same concept is further grounded in different neural network connections from one subject to the other. Such considerations offer new perspectives to apprehend language as a system of common shared cues, useful to make others live intended target experiences through the invocation of the right cues.

### 6.2 Further investigations

Further investigations must be conducted to challenge our theory or to further develop it.

#### Empirical methods

We verified in the literature the existence of retrograde signals that satisfy most of the assumptions of our theory. Further targeted empirical studies should verify the open ones so as to support (or invalidate) our claims. In particular, first, it is crucial to better study and validate the role of gap junctions in mediating backpropagating currents. Second, our literature review suggests that both glial gap junctions and electrical synapses seem to play a role in network oscillations related to memory recall. While mixed electrical and chemical synapses can create an environment in which chemical coupling is somewhat “copied” by electrical coupling (such that mixed neurons which are chemically coupled, are also electrically coupled), the role of glial cells in the entire process remains unclear.

#### Computational methods

Another line of work could study the computational effectiveness of our hypothesis in implementing its other target goals.

First, beyond object recognition and exact image reconstruction which we showcased in this work, less obvious tasks consist in reconstructing stimuli that has a temporal component such as speech or video. For this, neural network models that are more faithful to how humans process such temporal stimuli become much more paramount.

Another challenging task is imagination. There, a first sub-problem is how to “imagine” or reconstruct a plausible trace that corresponds to a generic concept, for example, imagining *any* cat, and not a particular *episodic* cat that we have previously seen. Once this first problem computationally modeled, then instead of backward-constructing only one concept at a time, we could simultaneously activate two concepts and observe the resulting reconstructed images. As exemplified before, one could activate a concept like “laughing” and a concept like “cat” and visualize the results of the “competition” between backpropagating signals on the backward-constructed images. This again will need neural networks that are not only richer than AI’s AlexNet in terms of features, but whose features exhibit certain biologically plausible arrangements (e.g. mechanisms similar to lateral inhibition or competitive neural processing whereby whenever a feature or perceptual interpretation is firing, the opposite alternative is suppressed). Of course, similar generative tasks can be achieved with today’s neural networks such as Dalle [231] but the latter lack biological plausibility [232, 233, 234].

Finally, another line of work should computationally model our theory in a more biologically plausible spiking neural network environment, taking into account more complex neuronal models where chemical and electrical synapses coexist.

## Footnotes

1 We discuss the existence of similar selective neurons in Sec. 3.3

2 The connection can be reinforced either by mere repetition, e.g. repeating many times the word cat in the presence of an image cat, or by neuromodulation, in which case a single co-occurrence can be enough.

3 one in theory should be enough but there must be many in reality, e.g. at least for redundancy to prevent failures

4 For simplicity, we focus in this toy example, on a single encounter that, we assume, was “encoded right away”. In reality, stored memory traces might evolve with repeated exposure, such that recalling what is meant by the word cat, leads to the recall of a memory of the signified, that is statistical in nature (e.g. one of the many cats met before, or an average abstract image of a cat)

5 The selectivity of pointer neurons is a consequence of learning to identify prior stimuli. This selectivity defines what particular contextual cues are efficient at recalling what trace.

6 A technique that allows to precisely reactivate or inhibit selected populations of neurons, that were previously tagged depending on their activity

7 On purpose, we use interchangeably encoding, learning and conditioning to accommodate terminologies of different papers.

8 While we give more attention in the remainder to electrical synapses, other types of gap junctions, such as those in glial cells or astrocytes, could potentially fulfill these criteria as well, perhaps complimentarily. Further research is needed to ascertain this.

9 “Connexins” refers to a family of proteins that form the channels making up gap junctions in vertebrate cells. These channels allow for direct communication of current and small molecules between the inside of adjacent cells.

10 Areas where sparse source pointer neurons have been found (See Sec 3.3).

11 While their involvement is well established, the details and functional split among them is still subject to continuous research [159].

12 This feature extends beyond mere action potentials as gap junctions enable the propagation of sub threshold activity between coupled neurons, suggesting a continuous[142], rather than discrete, mode of information transfer

13 often used to grossly approximate the spatial visual processing in the retina.

14 Unlike the original work which used a single split of train and test images, we performed different splits, sometimes reaching 99% accuracy and sometimes less.

15 Recent work [211] however has shown that training using “properly translated data” achieves the same effect as weight sharing.

16 In practice, we try different initializations [213] and pick the best

17 we simulate a single neuron for simplicity, but same reasoning applies multiple ones

18 In practice, implementation-wise, the score is simply the sum of weights. Indeed, the SNN model considers a spike to be a binary decision that happens when the accumulated potential reaches a threshold (i.e. we do not consider the rate). We tried a version that uses the exact value of the internal potential instead of the unit value 1 but results were similar.

19 Somewhat expected since the default learning rates were picked to maximize the performance of this particular 200/198 train/test split ratio.

20 Simple visual inspection of “good” and “bad” images did not uncover any peculiar character to distinguish them

21 This of course does not prove that this is how it happens in the humain brain. But it would be an additional argument in favour of the theory.

22 The model fails to reduce the loss below a certain value (a cross entropy of around 5) without overfitting to the training data.

23 Retraining all layers including *Conv*1 also yielded similar results, but there, we also inherited the “good-quality” weights of the original AlexNet.

24 Such lower level features are eventually shared by other objects and do not respond only to the presence of a cat.

## Notes

### Competing Interest Statement

The authors have declared no competing interest.

